# Zinc shapes the folding landscape of p53 and establishes a new pathway for reactivating structurally diverse p53 mutants

**DOI:** 10.1101/2020.07.23.217695

**Authors:** Adam R. Blanden, Xin Yu, Alan J. Blayney, Christopher Demas, Jeung-Hoi Ha, Yue Liu, Tracy Withers, Darren R. Carpizo, Stewart N. Loh

## Abstract

Missense mutations in the DNA binding domain (DBD) of the p53 tumor suppressor contribute to approximately half of new cancer cases each year worldwide. A primary goal in cancer therapy is to develop drugs that rescue the transcription function of mutant p53. Here we present a thermodynamic model that quantifies and links the major pathways by which mutations inactivate p53. The model is constructed by measuring folding free energies, zinc dissociation constants, and DNA dissociation constants of 20 of the most common DBD mutations in the p53 database. We report here that DBD possesses two unusual properties——one of the highest zinc binding affinities of any eukaryotic protein and extreme instability in the absence of zinc—which are predicted to cause p53 to be poised on the edge of folding/unfolding in the cell, with a major determinant being the concentration of available zinc. Eighty percent of the mutations examined impair either thermodynamic stability, zinc binding affinity, or both. Using a combination of biophysical experiments, cell based assays, and murine cancer models, we demonstrate for the first time that a synthetic zinc metallochaperone not only rescues mutants with decreased zinc affinities, but also mutants that destabilize DBD without impairing zinc binding. The latter is a broad class of p53 mutants of which only one member (Y220C) has been successfully targeted by small molecules. The results suggest that zinc metallochaperones have the capability to treat 120,500 patients per year in the U.S.

**SUMMARY:** Restoring tumor suppressing function to mutant p53 has the capability of treating millions of new cancer patients worldwide each year. An important step toward this goal is to categorize the spectrum of mutations based on how they inactivate p53. This study finds that the majority of the most common tumorigenic mutations compromise p53’s thermodynamic stability or its interaction with zinc, and demonstrates for the first time that members of both classes can be reactivated in cells by synthetic zinc metallochaperones. These results serve to stratify patients for potential zinc metallochaperone therapy.

## Introduction

The transcription factor p53 regulates a host of cellular responses to damage and distress ^1^. Its abilities to halt cell cycle progression, upregulate DNA repair pathways, and induce apoptosis help prevent deleterious mutations from propagating in cell populations. Mutations in p53 are an established driver of human cancer ^2^. The mutational spectrum of p53 is atypical because tumorigenic alterations are overwhelmingly missense and map to nearly every position within one of the domains of the protein (the DNA binding domain, or DBD) ^3^. By contrast, other frequently-mutated tumor suppressors such as BRCA1/2 ^4^, retinoblastoma 1 ^5^, and PTEN ^6^ are found with mostly nonsense, deletion, or insertion mutations. From a structural standpoint, p53 DBD is unusual in that it is marked by low thermodynamic and kinetic stability ^7^. The apparent melting temperature (T_m_) of wild-type (WT) DBD has been measured to be 32 – 45 °C depending on buffer composition ^8–10^, and the protein unfolds with a half-time of 9 m at 37 °C ^10^. As a result of this borderline stability, many tumorigenic mutations reduce T_m_ below body temperature and/or increase the rate at which the protein unfolds ^9,10^.

At basis of p53’s instability at physiologic conditions is its interaction with zinc. p53 consists of the N-terminal transactivation domain, a central DBD and the C-terminal tetramerization domain. The X-ray crystal structure of DBD (residues 94-312) reveals a β-sandwich with a DNA-binding surface consisting of a loop-sheet-helix motif and two loops (L2 and L3) (Fig. 1) ^11^. These loops are stabilized by the tetrahedral coordination of a single zinc ion by C176 and H179 of L2 and C238 and C242 of L3. Removing Zn^2+^ from DBD causes loss of DNA binding specificity, widespread changes in the protein NMR spectrum, and a reduction of ~3 kcal mol^−1^ in the apparent folding free energy ^12^. This inherent malleability has also been demonstrated in cells by overexpressing Zn^2+^-chelating proteins (metallothioneins) or adding small-molecule Zn^2+^ chelators, and observing reversible loss of sequence-specific DNA binding activity and a switch in recognition by an antibody that recognizes native p53 (PAB1620) to one that binds to unfolded/misfolded p53 (PAB240) ^13^.

**Figure 1.**
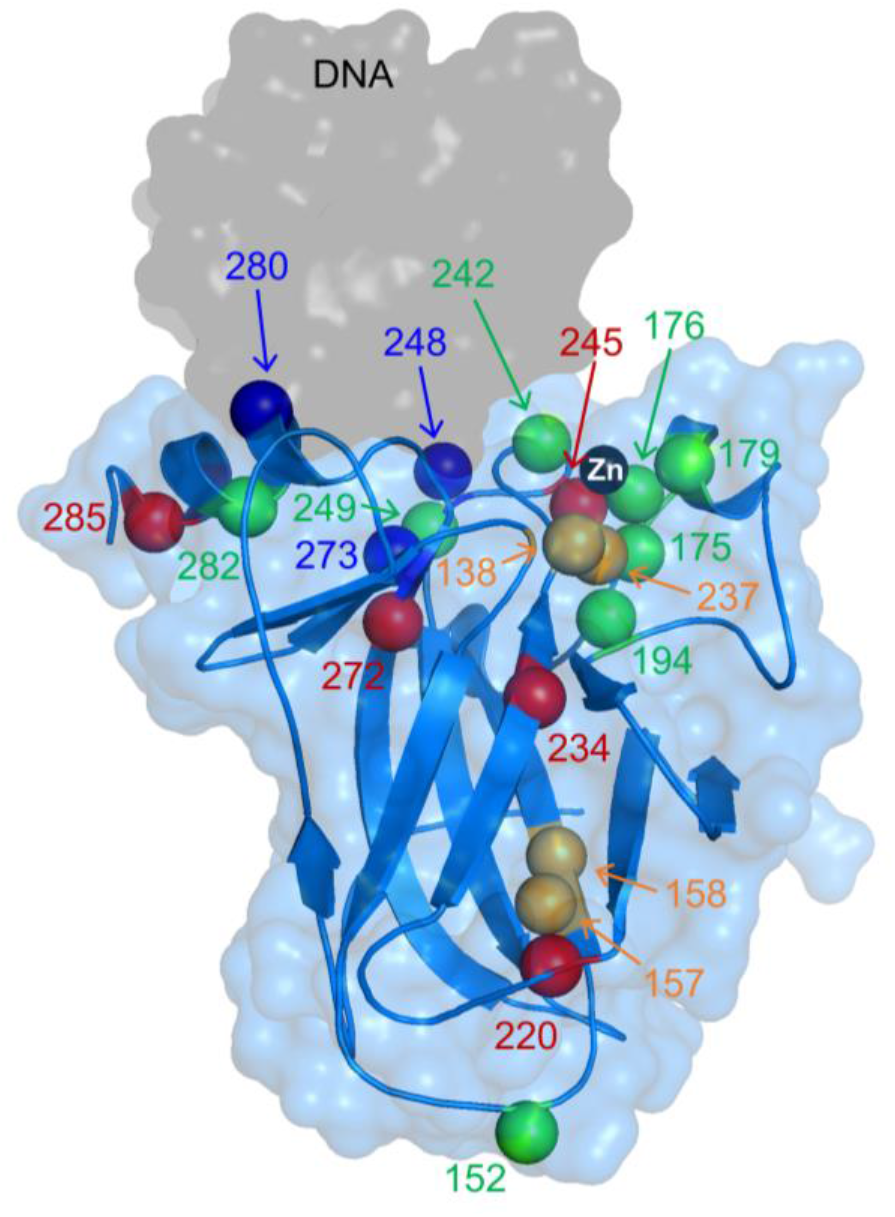
X-ray structure of WT DBD showing locations of mutations characterized in this study. Alpha carbons of mutated residues are colored according to their classifications described in *Results*: zinc-binding class (green), stability class (red), DNA-binding class (blue), and mixed zinc-binding/stability class (orange). DNA and Zn^2+^ are the gray surface and black sphere, respectively. PDB 1TSR.

Many tumorigenic mutations occur in and around the zinc binding site and presumably impair this interaction. The most common p53 mutation in cancer, R175H, is immediately adjacent to the zinc-chelating residue C176 (Fig. 1) and dramatically weakens metal binding affinity ^12,14^. Furthermore, there is evidence that some cancers inactivate p53 by upregulating metallothioneins and starving wild-type (WT) p53 of zinc ^15^. These observations have led our group and others to develop a new class of p53-targeted therapeutics based on Zn^2+^ delivery. By our definition, zinc metallochaperones (ZMCs) reactivate mutant p53 by shuttling Zn^2+^ from extracellular sources through the plasma membrane and into cells, thereby increasing intracellular Zn^2+^ concentrations to levels high enough to remetallate mutant p53. This approach has proven effective for multiple mutants in cell culture and mouse models of cancer ^14,16,17^.

Although the importance of zinc to p53 structure/function has long been recognized, there is still no quantitative thermodynamic model describing the p53-Zn^2+^ interaction and its linkage to folding. Previous attempts to characterize DBD folding thermodynamics have yielded valuable insight into mechanisms of p53 dysfunction, and have informed drug development efforts, but are ultimately incomplete because they lack information regarding zinc binding affinity of folded and unfolded states ^18,19^. Furthermore, there is disagreement in the literature regarding which mutants are potentially treatable using zinc-based therapies, likely because of the difficulty of inferring physical mechanisms from complex biological data and lack of a consensus definition regarding what constitutes a zinc-binding mutant ^17,20^.

Here we present and validate a thermodynamic model that partitions DBD folding energy into two measurable quantities: the free energy of folding of zinc-free DBD (apoDBD; ΔG_apo_), and the free energy of Zn^2+^ binding (ΔG_Zn_) to both native and non-native sites. The data reveal for the first time that: (i) ΔG_apo_ is extremely unfavorable at 37 °C, indicating that wild-type (WT) DBD is intrinsically unfolded in the absence of zinc, and (ii) DBD has one of the highest zinc binding affinities of any eukaryotic protein yet reported ^21^. Remarkably, the unusually large unfavorable value of ΔG_apo_ and the atypically large favorable value of ΔG_Zn_ are predicted to nearly cancel each other out at physiological temperature and intracellular zinc concentration, causing the overall free energy change of folding to be near zero.

Although p53’s instability at 37 °C has been previously documented, our modeling emphasizes that p53 is poised between folded and unfolded conformations in the cell, with available zinc concentration being a major determining factor. It further holds that mutations that decrease protein stability (but not zinc binding affinity) and mutations that decrease zinc binding affinity (but not protein stability) both cause p53 to lose function by a common unfolding mechanism, and both might be similarly rescued by increasing the concentration of available Zn^2+^. As a test, we apply the model to 22 of the most prevalent cancer-associated p53 variants and classify them into three classes—stability, zinc-binding, or DNA-binding—based on (respectively) ΔG_apo_, K_Zn_, and K_DNA_, the latter being dissociation constants for binding to a panel of p53 recognition elements. We validate the model by testing whether ZMC1 reactivates members of the stability class of p53 mutants in cells, a category not previously regarded as being amenable to Zn^2+^ therapy. The results provide a more complete picture of the p53 activation/inactivation landscape and significantly expand the number of p53 mutants that are potentially rescuable by ZMCs.

## Results

### DBD-zinc energy landscape

First, we developed a thermodynamic model to describe the conformational states of DBD as a function of [Zn^2+^]. We propose a minimal four-state mechanism in which Zn^2+^ binds to a single, high-affinity site in native apoDBD (N) with an equilibrium constant of 1/K_Zn_ (Eq. 1) and to one or more low-affinity sites in unfolded apoDBD (U) with an average equilibrium constant of 1/K_Zn,U_ (Eq. 2). HoloDBD (N·Zn^2+^) is the only species with native DNA binding activity.

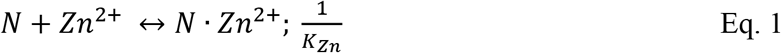

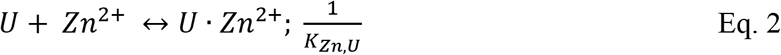

To measure K_Zn_, we monitored the increase in Tyr fluorescence ^12^ of apoDBD as it binds Zn^2+^ at 10 °C (Fig. 2A). [Zn^2+^]_free_ was buffered at the indicated concentrations using chelators of varying Zn^2+^ affinity. Fitting the data to the one-site binding equation yields K_Zn_ = (1.6 ± 0.3) × 10^−15^ M. To our knowledge, this is one of the lowest K_Zn_ values ever reported, with only one other eukaryotic protein possessing comparable affinity (PDZ and LIM domain protein 1; K_Zn_ = 3.2 × 10^−15^ M) ^21^. Because p53 is a tetramer, we measured K_Zn_ of the full-length protein to determine if there is cooperativity between the monomers (Fig. 2A). Fitting the data to the Hill binding equation reveals K_Zn_ = (0.4 ± 0.1) × 10^−15^ M and n = 0.99 ± 0.02, indicating that each monomer binds zinc independently and that the isolated DBD is an accurate representation of the DBD in the full-length p53 tetramer. We then measured K_Zn,U_ by competition assay between urea-denatured apoDBD and the fluorescent Zn^2+^ chelator FluoZin-3 (Fig. 2B). The apparent K_Zn,U_ value of (42 ± 7) × 10^−9^ M is 10^7^-fold weaker than K_Zn_. We generated a Ser mutant of the zinc-binding residue C176 and found its K_Zn,U_ to be essentially identical ((50 ± 23) × 10^−9^ M), indicating that the zinc binding sites in the native and unfolded proteins are distinct (Fig. 2B).

**Figure 2.**
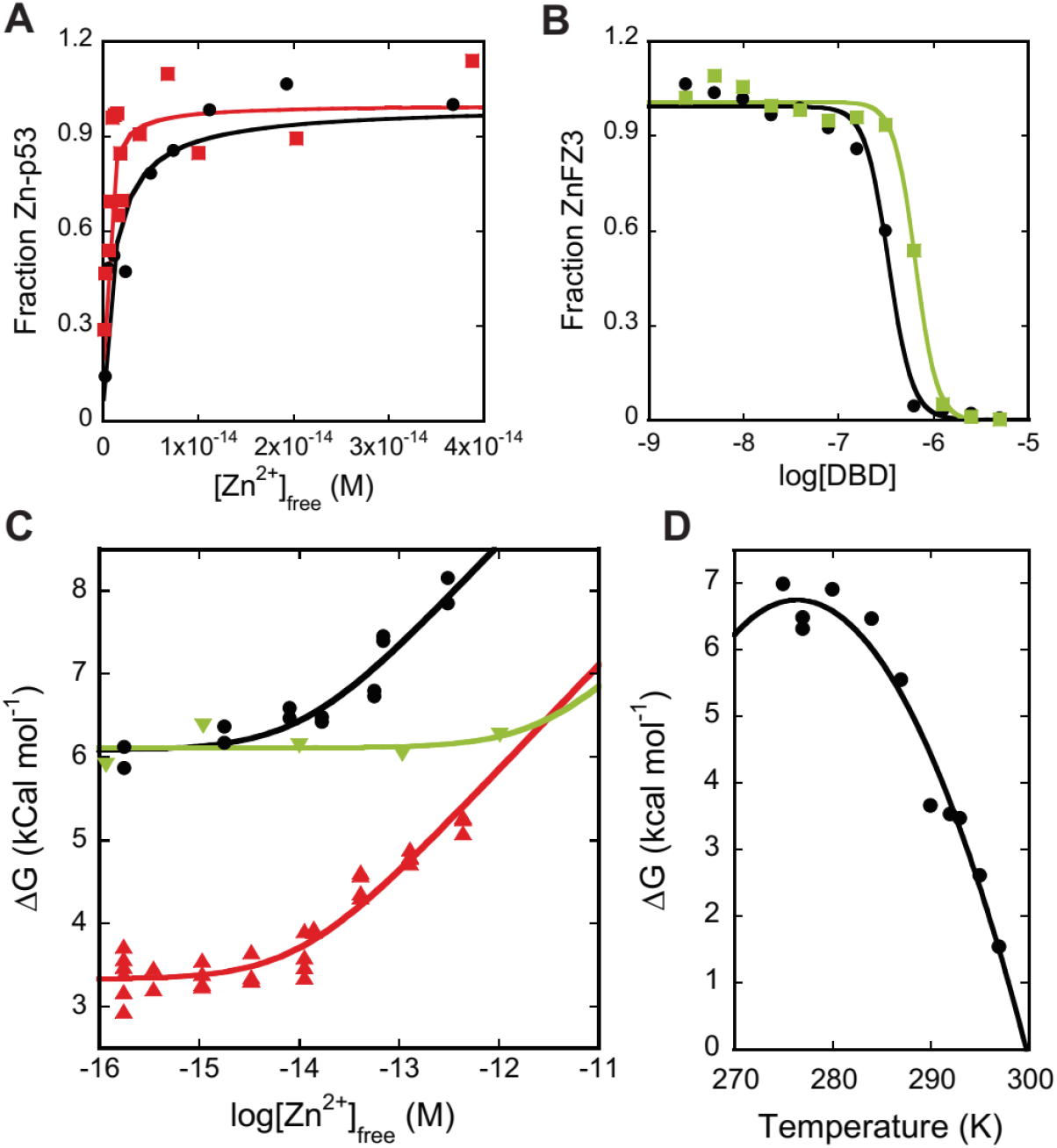
Zinc binding affinity and stability of DBD. **(A)** WT DBD (black) and full-length WT p53 (red) bind Zn^2+^ with K_Zn_ values of (1.6 ± 0.3) × 10^−15^ M and (0.4 ± 0.1) × 10^−15^ M, respectively, as determined by change in Tyr fluorescence (10 °C; n = 3). **(B)** Unfolded WT DBD (black) and unfolded C176S DBD (green), bind Zn^2+^ with K_Zn,U_ values of (42 ± 7) × 10^−9^ M and (50 ± 23) × 10^−9^ M, respectively, as determined by FluoZin-3 competition in 6 M urea (10 °C; n = 3). **(C)** Plotting folding free energy of DBD vs. [Zn^2+^]_free_ (10 °C) reveals that R175H (green) is a pure zinc binding-class mutant whereas A138V (red) is a pure stability-class mutant. The point at which the lines deflect upwards are the approximate K_Zn_ values. WT DBD is in black. **(D)** Temperature dependence of apoDBD folding free energy fit to the Gibbs-Helmholtz equation yields ΔH_m_ = 171 ± 20 kcal mol^−1^, T_m_ = 300 ± 1 K, and ΔC_p_ = 7.0 ± 1.7 kcal mol^−1^ K^−1^ (fit value ± SE of fit).

To delineate the relationship between zinc binding and WT DBD folding, we performed urea denaturation experiments at various concentrations of [Zn^2+^]_free_ that were fixed by a mixture of Zn^2+^ chelators (SI Fig. 1A). Zinc binding to unfolded DBD is negligible over the [Zn^2+^]_free_ range tested, so the overall free energy change for folding to the native state (ΔG_fold_) is given by Eq. 3, where ΔG_apo_ = −RT·ln(K_apo_) (Eq. 3) and ΔG_Zn_ = −RT·ln(K_Zn_)^−1^ (Eq. 1).

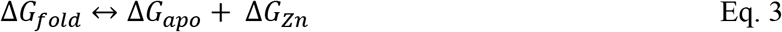

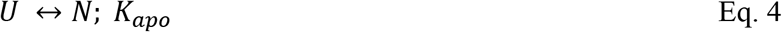

Zinc-induced stabilization can be visualized intuitively by plotting ΔG_fold_ against [Zn^2+^]_free_ (Fig. 2C). Focusing on the WT DBD data, the curve begins as a horizontal line that intersects the y-axis at ΔG_apo_, then starts to deflect upward when zinc begins to bind apoDBD, i.e. when [Zn^2+^]_free_ ≈ K_Zn_. ΔG_fold_ continues to increase linearly with log[Zn^2+^]_free_ and only levels off when zinc starts to bind unfolded DBD ([Zn^2+^]_free_ ≈ K_Zn,U_). A pure zinc-binding mutant such as R175H weakens ΔG_Zn_ (Eq. 1) without affecting ΔG_apo_, shifting the deflection point to higher [Zn^2+^]_free_ but maintaining the same y-intercept as WT (Fig. 2C). Conversely, the signature of pure stability mutants such as A138V, a well-established temperature sensitive variant ^22^ is a similar deflection point compared to WT but a y-intercept value closer to zero. We observed no trend in the cooperativity parameter (*m*-value) of the unfolding transitions over 10^−16^ M < [Zn^2+^]_free_ < 10^−12^ M (SI Fig. 1A), signifying that urea denaturation is adequately described by a two-state transition between unfolded and native apo or native holo states (the m-values of the two are similar ^12^) throughout the tested zinc concentration. We also performed denaturation experiments starting with either apoDBD or zinc-bound DBD and obtained similar results, indicating that metal binding had reached equilibrium during the incubation time (SI Fig. 1B). The data are described well by Eq. 3 and yield fit parameters of ΔG_apo_ = 6.4 ± 0.1 kcal mol^−1^ and K_Zn_ = (7.0 ± 2.5) × 10^−15^ M, both of which are in good agreement with direct measurements (Table 1).

**Table 1:** Stabilities and zinc binding affinities of apoDBD variants (10 °C). ΔG_apo_ and K_Zn_ were obtained from bi-linear fits of the data in SI Fig. 2 (errors are SE of the fits). K_Zn_ (direct) values were obtained from Tyr fluorescence spectra as in Fig. 2A (errors are SD, n = 3).

| **Table 1: Stabilities and zinc binding affinities of apoDBD variants (10 °C).** $\Delta G_{apo}$ and $K_{Zn}$ were obtained from bi-linear fits of the data in SI Fig. 2 (errors are SE of the fits). $K_{Zn}$ (direct) values were obtained from Tyr fluorescence spectra as in Fig. 2A (errors are SD, n = 3). | | | |
| --- | --- | --- | --- |
| Mutant | $\Delta G_{apo}$ (kcal mol<sup>-1</sup>) | $K_{Zn}$ (M) | $K_{Zn}$ (direct) (M) |
| WT | $6.3 \pm 0.1$ | $7 \pm 2 \times 10^{-15}$ | $1.7 \pm 0.7 \times 10^{-15}$ |
| **Zinc binding class** |  |  |  |
| C176S | $5.9 \pm 0.1$ | | $2.7 \pm 0.8 \times 10^{-10}$ |
| R175H | $6.3 \pm 0.2$ | | $5.2 \pm 1.8 \times 10^{-11}$ |
| C242S | $5.8 \pm 0.1$ | | $2.1 \pm 0.6 \times 10^{-12}$ |
| L194F | $7.17 \pm 0.06$ | $5.0 \pm 1.9 \times 10^{-14}$ | |
| R249S | $6.0 \pm 0.1$ | $3.3 \pm 1.8 \times 10^{-13}$ | |
| H179R | $6.3 \pm 0.1$ | | $1.1 \pm 0.5 \times 10^{-13}$ |
| P152L | $6.2 \pm 0.3$ | $5.4 \pm 6.6 \times 10^{-14}$ | |
| R282Q | $5.8 \pm 0.2$ | $3.3 \pm 1.9 \times 10^{-14}$ | |
| S241F | $5.8 \pm 0.2$ | $3.1 \pm 1.7 \times 10^{-14}$ | |
| **Stability class** |  |  |  |
| E285K | $1.7 \pm 0.1$ | $1.0 \pm 0.3 \times 10^{-14}$ | |
| Y234C | $3.3 \pm 0.1$ | | $1.9 \pm 0.2 \times 10^{-15}$ |
| Y220C | $4.2 \pm 0.1$ | $2.6 \pm 1.9 \times 10^{-14}$ | |
| V272M | $4.30 \pm 0.06$ | $2.3 \pm 0.4 \times 10^{-15}$ | |
| A138V | $3.33 \pm 0.05$ | $1.0 \pm 0.2 \times 10^{-14}$ | |
| G245S | $5.3 \pm 0.1$ | $3.5 \pm 1.7 \times 10^{-14}$ | $7.1 \pm 2.5 \times 10^{-15}$ |
| **Mixed zinc binding/stability class** |  |  |  |
| V157F | $3.4 \pm 0.2$ | $9.5 \pm 8.5 \times 10^{-14}$ | |
| R158H | $5.0 \pm 0.2$ | $5.1 \pm 3.6 \times 10^{-14}$ | |
| M237I | $5.2 \pm 0.2$ | | $3.3 \pm 1.5 \times 10^{-13}$ |
| **DNA binding class** |  |  |  |
| R248Q | $7.6 \pm 0.5$ | $6.5 \pm 9 \times 10^{-15}$ | $1.0 \pm 0.2 \times 10^{-15}$ |
| R273H | $7.3 \pm 0.2$ | $1.4 \pm 0.9 \times 10^{-14}$ | |
| R280K | $7.89 \pm 0.08$ | $1.6 \pm 0.9 \times 10^{-14}$ | |

### Extrapolation to physiological conditions

Because our DBD experiments must be performed at low temperature to avoid protein aggregation, we sought to gain insight into how ΔG_apo_ changes with temperature. To quantify this, we performed urea melts of apoDBD at 10 temperatures over the range 2 °C – 22 °C. The resulting curve is a parabola the narrowness of which is proportional to ΔC_p_, the change in heat capacity on protein unfolding at constant pressure (Fig. 2D). We fit these data to the Gibbs-Helmholtz equation (SI Fig. 1C) to obtain the heat capacity change of unfolding (ΔC_p_), T_m_, and the enthalpy change of unfolding at T_m_ (ΔH_m_). ΔC_p_ of apoDBD (7.0 ± 1.7 kcal mol^−1^ K^−1^) is substantially higher than the value previously calculated for holoDBD (3.8 kcal mol^−1^ K^−1^) ^18^ as well as that predicted based on the size of DBD and change in accessible surface area upon unfolding ^23^. To eliminate the possibility of buffer or denaturant-specific effects, we changed the buffer from Tris to phosphate and the denaturant from urea to guanidine hydrochloride and obtained nearly identical results (SI Fig. 1D). There was no trend in *m*-value that would signify deviation from two-state behavior (SI Fig. 1E). We therefore conclude that apoDBD, owing to its large ΔC_p_ value, possesses a stability that is unusually dependent on temperature.

We then combined the full four-state model (Eq. 1, Eq. 2, Eq. 4) with the Gibbs-Helmholtz equation to project the energy landscape of WT DBD as a three-dimensional plot with [Zn^2+^]_free_, temperature, and fraction holoDBD (F_holo_) on the x-, y-, and z-axes, respectively (Fig. 3A). When the protein is kept cool (10 °C) it is stable even without zinc (ΔG_apo_ = −6.3 kcal mol^−1^), making F_holo_ effectively unity at all relevant concentrations of free zinc (> 10^−15^ M). At 37 °C, however, the protein is completely unfolded without zinc (ΔG_apo_ = 6.9 kcal mol^−1^). Strikingly, the high zinc binding affinity of WT apoDBD together with the concentration of free zinc in the typical cell (~10^−10^ M) ^24,25^ results in an overall ΔG_fold_ of ~0 kcal mol^−1^ and F_holo_ ~0.5 (Fig. 3A). This suggests that under normal physiological conditions, WT p53 is balanced on the edge of folding and unfolding and may be pushed in either direction by surprisingly small changes in [Zn^2+^]_free_, ΔG_apo_, K_Zn_, or temperature. For example, raising [Zn^2+^]_free_ to 10^−9^ M or lowering it to 10^−11^ M yields F_holo_ values of 0.9 and 0.08, respectively. F_holo_ changes similarly when K_Zn_ is altered by a factor of 10, ΔG_apo_ by 1-2 kcal mol^−1^, and temperature by 2 °C. The energy landscape of mutant DBD can be generated in the same manner as that of WT DBD provided that the mutation does not alter the enthalpy change of unfolding. This scenario is reasonable for the zinc-binding class, because these mutations do not perturb ΔG_apo_ and thus ΔH and ΔS can be assumed to remain unchanged. Stability-class mutations, however, can affect ΔH, ΔS, or both, making the extrapolation of ΔG_fold_ to 37 °C unreliable.

**Figure 3.**
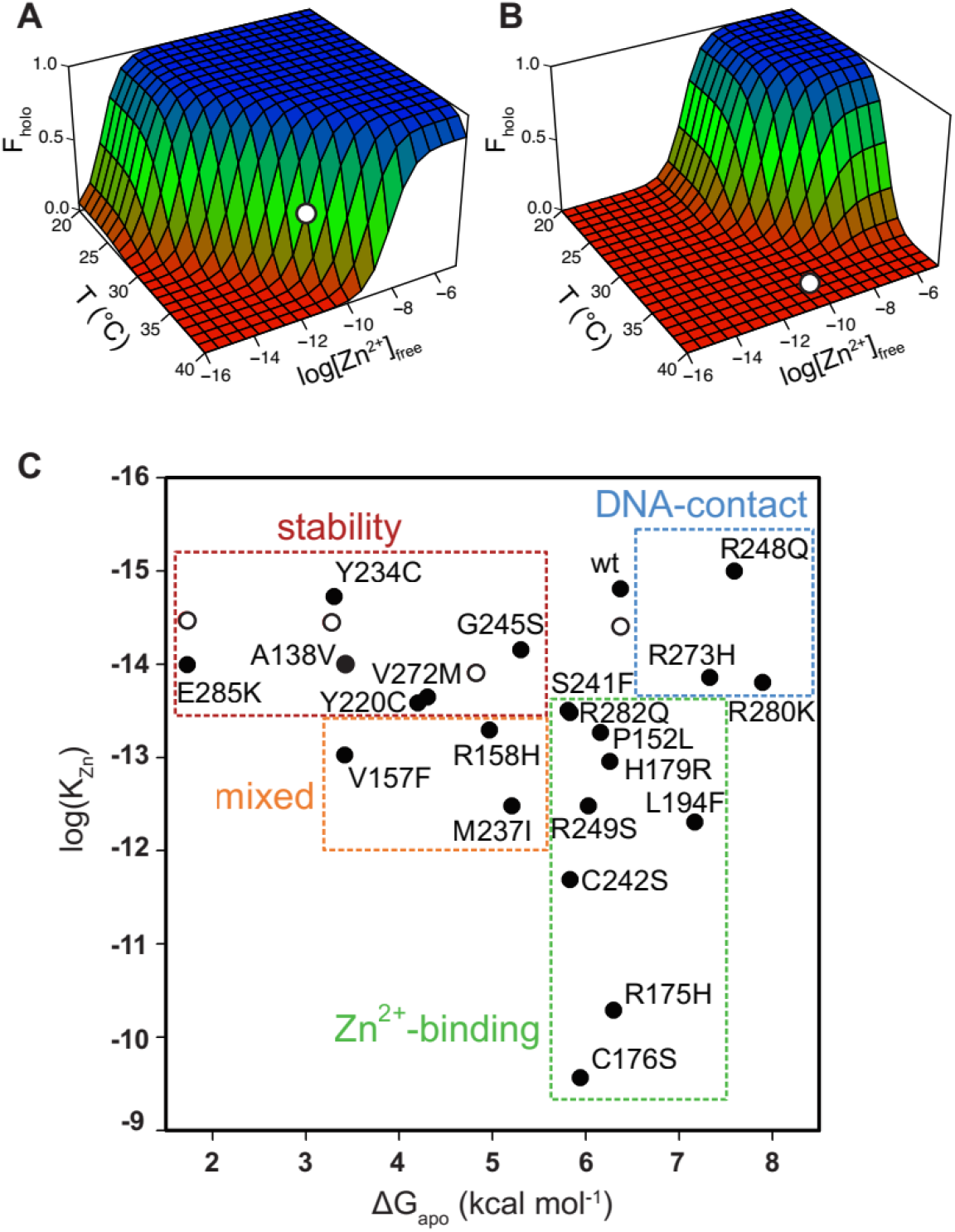
Energy landscape of DBD folding and classification of p53 mutants. The populations of folded, active WT DBD **(A)** and R175H DBD **(B)** depend strongly on free zinc concentration and temperature. White circles indicate physiological T and [Zn^2+^]_free_. **(C)** 17 of the top 20 most common tumorigenic p53 mutations impair DBD thermodynamic stability (red box), zinc binding affinity (green box), or both (orange box) (10 °C). The remaining three are DNA contact mutations. Open circles indicate WT DBD destabilized by urea.

### Categorization of p53 mutants

We then sought to apply this methodology to gain insight into the mechanisms by which tumorigenic mutations cause p53 to lose function, we purified 22 DBD missense mutants commonly found in human cancer. We chose the most frequent somatic mutations in the IARC database with the following additional criteria: (i) when multiple mutations were reported at a single position, only the most common variant was used (to maximize protein coverage); (ii) if the residue was mutated to Trp, we selected the next most frequent alteration at that position (an extra Trp interferes with our fluorescence assays); (iii) when multiple mutations of a zinc-coordinating residue were common, we chose the most isosteric in order to help isolate the effects of Zn^2+^ binding. We performed urea denaturation experiments as a function of buffered [Zn^2+^]_free_ to determine ΔG_apo_ and K_Zn_ at 10 °C (SI Fig. 2), and we also cross-checked K_Zn_ using direct Zn^2+^ binding experiments (c.f. Fig. 2A) for select mutants. We were able to determine ΔG_apo_ for all mutants and K_Zn_ for all but two (Y205C, Y163C; *vide infra*) (Table 1).

By placing the 22 variants on a plot according to their ΔG_apo_ and K_Zn_ values, three classes of mutants emerge: pure zinc-binding, pure stability, and DNA-binding, with a sub-class of mixed zinc-binding/stability phenotype (Fig. 3C). Pure zinc-binding mutants exhibit decreased Zn^2+^ affinity but normal stability and thus cluster near a vertical line drawn directly below WT DBD (green box). Members of this class include R175H as well as the direct Zn^2+^ ligating mutants C176S, C242S, and H179R. Significantly, the analysis reveals a number zinc-binding mutations (P152L, L194F, R282Q) that were not previously suspected as being so due to their long distances from the metal binding pocket (Fig. 1).

Pure stability mutants are less stable than WT but suffer no zinc-binding deficiency; these are defined by falling near the horizontal line drawn to the left of WT DBD (red box in Fig. 3C). To validate that protein stability can indeed be separated from zinc affinity in this manner, we destabilized WT apoDBD using sub-denaturing amounts of urea and measured K_Zn_ using the Tyr fluorescence assay. These data (open circles) fall on the horizontal line as predicted by the model. The mixed-phenotype mutants (A138V, V267F, R158H, M237I; orange box in Fig. 3C) decrease both stability and zinc affinity. Only three mutations (R248Q, R273H, R280K) do not impair either stability or metal affinity (blue box in Fig. 3C). These residues are in direct contact with DNA (Fig. 1) and the mutants were known to possess stabilites greater than or equal to that of than WT ^12,18^. Our data confirm that they bind zinc normally.

Figure 3C reveals a remarkable range in magnitude of the stability and zinc binding deficiencies. The most severe member of each class reduces ΔG_apo_ by 4.6 kcal mol^−1^ (E285K) and K_Zn_ by 5 orders of magnitude (C176S). Additionally, several zinc-binding or mixed mutations (L194F, R282Q, V157F, R158H, P152L) lie far from the canonical zinc-binding pocket, suggesting long-range communication between distant regions of DBD. The subtle nature of DBD structure and energetics is also evident from the observation that adjacent mutations can produce markedly different effects. For example, R248Q is a classic DNA contact mutant whereas R249S is a pure zinc-binding mutant. This demonstrates the difficulty in inferring physical consequences of p53 mutation based on location and stresses the need to measure properties of each mutant to gain a clear understanding of its impairment.

### DNA binding affinity

To gain a better understanding of the effects of p53 mutation on DNA binding activity, we measured dissociation constants (K_DNA_) for binding of the 22 DBD mutants to a panel of fluorescently labeled oligonucleotides bearing 10 p53 recognition elements (p53RE; SI Table 1). The fluorescence anisotropy data fit adequately to the Hill binding equation. When global fits for individual sequences were performed by linking the Hill parameter and curve amplitudes, the resultant curves fit equally well regardless of DBD mutant, indicating similar binding mechanisms regardless of mutation (SI Fig. 3A). SI Figure 3B represents the K_DNA_ values relative to that of WT DBD in the form of a heat map. We excluded the two p53REs to which none of the DBD variants bound (WAF1-3’ and EGFR), and assigned K_DNA_ = 25 μM (the weakest value we could consistently measure) to data sets in which we could detect no interaction.

By visual inspection, most of the stability and mixed mutants maintain similar binding affinity for the different p53REs, whereas all of the DNA-contact mutants and most of the zinc-binding mutants lose measurable affinity for all p53REs (SI Fig. 3B). The majority of zinc-binding mutants purify with sub-stoichiometric but detectable zinc content (e.g. R175H contains 0.6 equivalents of Zn^2+ 14^), and consequently should show partial DNA binding activity. We speculated that the loss of DNA binding may be caused by Zn^2+^ misligating to non-native sites (K_Zn,U_ = 42 × 10^−9^ M; Fig. 2B) that out-compete the native site at the temperature of protein expression (18 °C). To test this hypothesis, we attempted to re-establish native metallation status to the zinc-binding mutants P152L, R175H, H179R, and L194F by first removing all bound metal and then remetallating using a EGTA/ZnCl_2_ buffering system. This procedure restored measurable DNA binding affinity (WAF1 oligonucleotide) to all four mutants (SI Fig. 3C). The positive control (WT) and negative control (R280K) yielded the expected results of normal and undetected DNA binding, respectively. These data suggest that zinc misligation contributes to loss of DNA binding activity for this class of mutants.

Interestingly, while most of the data are explained by global increases and decreases in affinity based on thermodynamic category, there do seem to be mutation-specific effects as well. P152L maintains WT-like affinity for WAF1 5’, GADD45, and RGC, but loses all measurable affinity for PUMA, Type IV collagenase, p53RFP, MDM2, and BAX. As another example from the stability mutant category, Y220C maintains affinity for WAF1 5’, GADD45, PUMA, and Type IV Collagenase, but seems to lose affinity for p53RFP, MDM2, BAX, and RGC. Additionally, E285K gains affinity for all p53REs, but the increase in affinity ranges from 3-fold for WAF1 5’ to 15-fold for MDM2. E285K may bind more tightly to all DNA sequences due to enhanced electrostatic interactions with the phosphate backbone afforded by the negative-to-positive charge reversal near the active site. These results indicate that there are conserved functional consequences of the thermodynamic impairments we measure for p53 mutants that account for the majority of the functional differences we see in DNA-binding phenotype, but there are also mutation-specific effects that are better explained by idiosyncratic structural changes caused by each mutant that our model does not capture.

### Tyr to Cys mutants are exceptions to the model

Y163C, Y205C, Y220C, and Y234C are all destabilized (ΔG_apo_ = 3.5 – 5.0 kcal/mol), but, unlike all other members of the stability class except for M237I, their ΔG_fold_ values fail to increase with [Zn^2+^]_free_ (SI Fig. 2). The flat profiles of these plots can potentially be explained by very weak K_Zn_ values, but two observations argue against this interpretation. First, direct measurement of K_Zn_ by Tyr fluorescence yields a value of (1.9 ± 0.2) × 10^−15^ M for Y234C (Table 1), the only Tyr→Cys mutant that was able to survive the zinc removal procedure without aggregating. Second, Y205C, Y220C, and Y234C retain WT-like DNA binding activity, whereas most of the zinc-binding mutants are compromised in this regard. We hypothesized that the extra Cys residue forms a new Zn^2+^ interaction site in the unfolded state, thereby bringing K_Zn,U_ closer to K_Zn_ and siphoning the native state of the stabilization energy it would normally receive by binding Zn^2+^. To test this hypothesis, we denatured Y234C apoDBD in 6 M urea and measured K_Zn,U_ by Tyr fluorescence. Unfolded Y234C binds zinc twice as tightly as WT [K_Zn,U_ = (20 ± 4) × 10^−9^ M; n=6], consistent with the hypothesis with the caveat that the unfolded state in urea is likely different from that in buffer. As an additional test, we made the Y234A mutant and found that this mutation restores the relationship between ΔG_fold_ and [Zn^2+^]_free_ as predicted by the model (SI Fig. 2), with a K_Zn_ value identical to that of WT within experimental error (Table 1). These results demonstrate that Y234A is a stability-class mutant and suggest that other Tyr→Cys mutants may also be members of this category. The data also indicate that introducing Cys can encourage zinc misligation in the unfolded state and destabilize native p53 through manipulation of K_Zn,U_ in our model.

### Testing the energy landscape model in cells using ZMC1

We previously demonstrated that the small-molecule zinc metallochaperone ZMC1 can reactivate p53 mutants of the zinc-binding class in cultured cells. ZMC1 forms a 2:1 complex with Zn^2+^ and acts as an ionophore to transport Zn^2+^ into the cell, whereupon it buffers [Zn^2+^]_free_ to 10 – 20 nM ^14^. When ZMC1 was added to human cancer cell lines homozygous for p53^R175H^ or one of the direct zinc ligation mutations (p53^C176F^, p53^C238S^, p53^C242S^), cell toxicity was observed with EC_50_ values well below that of p53^WT^ and p53-null controls. This enhanced ZMC1 sensitivity was shown to be due to a p53-mediated apoptotic program ^26^. Given the ZMC mechanism, we previously surmised that the spectrum of mutants to which ZMCs were amenable was limited to those with impaired zinc binding. However, the relationship of zinc binding to the energy of protein folding we observed in our model allowed us to hypothesize that raising intracellular zinc concentrations could increase protein stability enough to rescue wild type conformation of some stability mutants. To test this, we used ZMC1 as a tool to raise intracellular concentrations of zinc in an array of zinc-binding, stability, and mixed classes of p53 mutants generated by site specific mutation in plasmids and expressed in H1299 (p53-null) cells. We also used human tumor cell lines that endogenously express mutant p53, when available. Cell survival curves are shown in SI Fig. 4B and ZMC1 EC_50_ values are summarized in Table 2.

**Table 2.** ZMC1-mediated toxicity and p53 refolding in H1299 cells expressing p53 mutants.

| **Mutant** | **EC<sub>50</sub> (nM)** | **p53 refolding<sup>a</sup>** |
| --- | --- | --- |
| Untransfected cells | > 1000 | Not applic. |
| Empty vector | > 1000 | Not applic. |
| **Zinc-binding class** |  |  |
| R175H | 5 | Yes |
| C176S | 2 | Yes |
| L194F | 5 | Yes |
| P152L | > 1000 | No |
| R282Q | > 1000 | No |
| **Stability class** |  |  |
| Y234A | 10 | Yes |
| Y234C | > 1000 | No |
| V272M | 2 | Yes |
| E285K | > 1000 | No |
| **Mixed zinc-binding/stability class** |  |  |
| M237I | > 1000 | Yes |
| **DNA binding class** |  |  |
| R273H | > 1000 | Yes |
| R280K | > 1000 | No |
| <sup>a</sup>Yes, PAB240 staining level was reduced significantly (Fig. 4A) after ZMC1 treatment; No, PAB240 staining level was not significantly changed after ZMC1 treatment. |  |  |

The mutants tested in the pure zinc-binding class are R175H, C176S, L194F, P152L, and R282Q. Cells transfected with R175H, C176S, and L194F show >100-fold enhanced sensitivity to ZMC1 (EC_50_ = 0.002 μM – 0.005 μM) relative to untransfected and vector-only controls (Table 2). A p53^L194F^ tumor cell line is also sensitive to ZMC1 (SI Fig. 5A). EC_50_ values of R282Q and P152L are not detectable in the tested range, suggesting that they are not functionally reactivated by ZMC1. The negative control group consists of the DNA-contact mutants R273H and R280K. As expected, these variants show no increased sensitivity to ZMC1, indicating that their DNA binding defects cannot be ameliorated in a zinc-dependent fashion.

The most discriminating test of the thermodynamic model is whether the pure stability-class (Y234C, Y234A, V272M, E285K) and mixed stability/zinc-binding class (M237I) of p53 mutants can be reactivated by elevating intracellular zinc. V272M is marked by pronounced ZMC1 sensitivity, with an EC_50_ value (0.002 μM) lower than that observed for any of the zinc-binding class mutants. This result demonstrates that a mutation whose sole consequence is to promote unfolding of p53 can be rescued by zinc. Y234C, E285K, and M237I, however, fail to show ZMC1 sensitivity. The negative result for Y234C agrees with the biophysical data (SI Fig. 2), which suggested that the extra Cys residue forms a competing, non-native zinc binding site in the unfolded state. We tested that hypothesis using the Y234A mutant. As predicted by the zinc misligation model, the EC_50_ value of Y234A drops to a value comparable to that of the pure zinc-binding mutants R175H and L194F (Table 2). To confirm the result for M237I, we tested two human cancer cell lines that express p53^M237I^ and found that they too were insensitive to ZMC1 (SI Fig. 5B). A human cancer cell line bearing p53^Y220C^ also showed insensitivity to ZMC1 (SI Fig. 5B).

### p53 refolding in cells monitored by conformation-specific antibody

To further test the thermodynamic model, and to gain insight as to why some mutants fail to regain their cell-killing functions in the presence of ZMC1, we assayed the extent to which elevated zinc can refold mutant p53 in cells. Refolding was monitored by the reduction in staining by the antibody PAB240, which binds to a cryptic epitope only exposed on p53 unfolding or misfolding (residues 212 – 217 ^27^). For the zinc-binding class of mutants, the variants that exhibit low EC_50_s (R175H, C176S, L194F) show reduced binding to PAB240 after ZMC1 treatment (Fig. 4A and SI Fig. 6A), implying that they undergo zinc-mediated refolding. The human cancer cell line T47D bearing the p53^L194F^ mutation also showed refolding of mutant p53 by reducing PAB240 staining and increasing PAB1620 staining (SI Fig. 5C). Antibody staining of P152L and R282Q did not change after ZMC1 treatment, consistent with their lack of sensitivity to the drug (Fig. 4A). It is possible that they misfold and/or aggregate in the cell to a conformation that is not amenable to zinc binding or refolding. As expected, staining intensities of the DNA-contact mutants R273H and R280K do not change after ZMC1 treatment. R280K cells appear dark in the absence of ZMC1, consistent with the DNA-contact phenotype of this mutant, but R273H stains brightly. This result suggests that the R273H mutation induces misfolding or aggregation in addition to loss of DNA binding affinity by direct contact.

**Figure 4.**
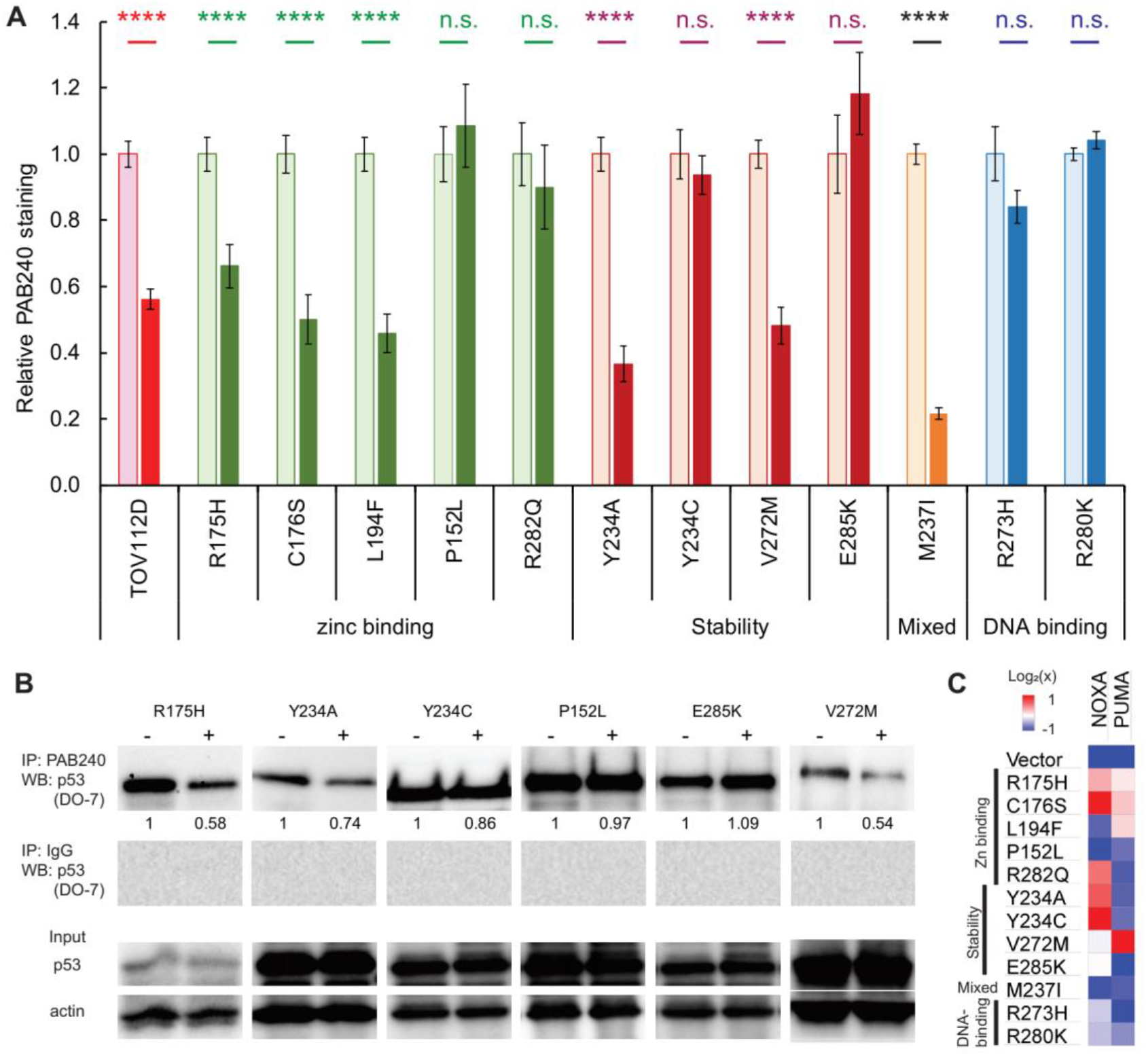
Response of p53 mutants to ZMC1 treatment in cells. **(A)** ZMC1-induced folding of p53 mutants quantified by PAB240 immunofluorescence. H1299 cells were stably transfected with p53 mutants and treated with 1 μM ZMC1 (dark bars) or DMSO vehicle (light bars) for 4 h. TOV112D (p53^R175H^) cancer cell line and parental cell line are positive and negative controls, respectively. ****, p < 0.0001; n.s., not significant. **(B)** ZMC1-induced folding of p53 mutants quantified by PAB240 IP. Protein lysates were extracted from cells, immunoprecipitated with PAB240, and blotted with the pan-p53 antibody DO-7. **(C)** Activation of *PUMA* and *NOXA* expression by ZMC1 (1 μM, 24 h) in stably transfected H1299 cells quantified by RT-PCR. Color shades indicate 2-fold differences relative to the vehicle-only controls.

For the pure stability-class mutants, the PAB240 refolding results (Fig. 4A) agree well with the EC_50_ data (Table 2): the mutants that regain cell-killing activity in the presence of ZMC1 (Y234A and V272M) show reduced PAB240 antibody staining, and the mutants that are insensitive to ZMC1 (Y234C and E285K in H1299 cells and Y220C in a cancer cell line (SI Fig. 5B)) react equally well with PAB240 before and after drug treatment. The mixed stability/zinc-binding mutant M237I is an interesting exception. M237I shows the greatest decrease in PAB240 staining after ZMC1 treatment of all mutants tested, yet M237I cells are insensitive to the drug. Increasing intracellular zinc therefore appears to refold M237I, as predicted by our model, but fails to restore its apoptotic activity. In agreement, the human cancer cell lines with p53^M237I^ mutation also showed refolding after treatment (by decreasing PAB240 staining after treatment of ZMC1) (SI Fig. 5E) but was not sensitive to ZMC1 (SI Fig. 5B). The M237I mutation may compromise p53 function by an additional mechanism such as introducing a structural defect in the folded protein or perturbing a binding interaction between p53 and another protein. We further tested the DNA binding of p53^M237I^ to the p53RE in p21 promoter in cells using a luciferase reporter assay and found that the M237I mutant was unable to activate transcription (SI Fig. 7).

As immunofluorescence is sometimes criticized for its possible interference with protein structure by fixation of cells, we sought to confirm refolding of the p53 mutants by performing immunoprecipitation using PAB240 followed by western blot for p53 proteins. Consistent with the immunofluorescent staining of the individual cells, R175H, Y234A, V272M showed decreased p53 protein by PAB240 pull-down after ZMC1 treatment, while the p53 band intensities remained similar for Y234C, P152L and E285K (Fig. 4B). As a positive control, we treated the temperature-sensitive A138V with ZMC1 and observed a 2-fold decrease in PAB240 pull-down (SI Fig. 8).

### Restoration of p53 transcriptional function

To determine if the conformational change observed with the mutants results in restoration of WT p53 transcriptional function, we compared mRNA levels of the p53-responsive genes *PUMA* and *NOXA* before and after 24 h of ZMC1 treatment, using the transfected cells described above. Mutants for which no increase in transcription is observed for either of the two probe genes (blue or white bars in Fig. 4C) are P152L and E285K (stability class), M237I (mixed class), and R273H (DNA-contact class). All these mutants are insensitive to ZMC1-mediated cell killing and fail to refold as judged by the PAB240 test (except M237I). Mutants for which ZMC1 treatment enhances transcription of at least one gene (one or more beige or red bars in Fig. 4C) are R175H, C176S, L194F, R282Q (zinc-binding class), Y234A, Y234C, and V272M (stability class), and R280K (DNA contact class). Thus, all the mutants that exhibit low EC_50_ values and undergo ZMC1-dependent refolding also have their transcriptional activities at least partially restored by the drug. Similarly, the human cancer cell line with p53^L194F^ also showed decreased mutant p53 protein (SI Fig. 5C) and the induction of the gene expression of *p21* and *PUMA* (SI Fig. 5E), indicating reactivation of p53 transcriptional function. The transcriptional assay, however, also finds that ZMC1 elevates mRNA levels of at least one gene for two stability mutants (Y234C and R282Q; Fig. 4C) that failed to show enhanced cell killing or refolding in the presence of ZMC1. The increase in mRNA levels appears to be insufficient to bring about apoptosis.

### V272M and E285K stability mutants

The stability mutant V272M is a particularly interesting case because it is efficiently refolded by ZMC1 (Fig. 4A) and is three-fold more sensitive to the drug than the pure zinc-binding mutant R175H (Table 2). This behavior is consistent with the thermodynamic model. The energy landscape of R175H indicates that raising intracellular [Zn^2+^]_free_ from 100 pM to 100 nM increases the percentage of folded R175H from 0.0028 % to 2.8 % (Fig. 3B). V272M is destabilized by 2.0 kcal mol^−1^ and if we assume that this relatively small perturbation does not change the enthalpy of unfolding, then V272M will achieve 2.8 % refolding at only 10 – 20 nM [Zn^2+^]_free_. Both IF and IP confirm ZMC1 refolds V272M, whereas ZMC1 fails to induce refolding of the negative control (R273H) (Fig. 4A; Fig. 4B). Finally, ZMC1 elicits high level of *PUMA* and *NOXA* expression in V272M (Fig. 4C). These findings demonstrate that V272M is robustly activated by ZMC1.

E285K, the most severe of the stability class of mutants, failed to refold in the presence of ZMC1 (Fig. 4A; Fig. 4B) and cells transfected with *p53*^E285K^ were not sensitive to the drug (Table 2). We speculated that the extreme instability of E285K prevented it from refolding even at the elevated intracellular zinc concentrations afforded by ZMC1 treatment. We asked whether reducing temperature could act synergistically with ZMC1 to reactivate E285K. The temperature of p53^E285K^ H1299 cell cultures was lowered to 22 °C for 4 h in the presence of ZMC1 to allow for temperature-assisted refolding, then returned to 37 °C for cell-killing assays. This procedure resulted in 3.8-fold decrease in EC_50_ compared to the control in which 37 °C was maintained throughout (Fig. 5A, Fig. 5B). Of note, we observed a similar effect for R175H in H1299 (2.4-fold decrease) and TOV112D cells (1.6-fold decrease), suggesting that low temperature acts synergistically with zinc-binding and stability mutants alike, as predicted by the thermodynamic model. We further determined that reducing temperature to 22 °C successfully induced ZMC1-mediated refolding of E285K (Fig. 5C).

**Figure 5.**
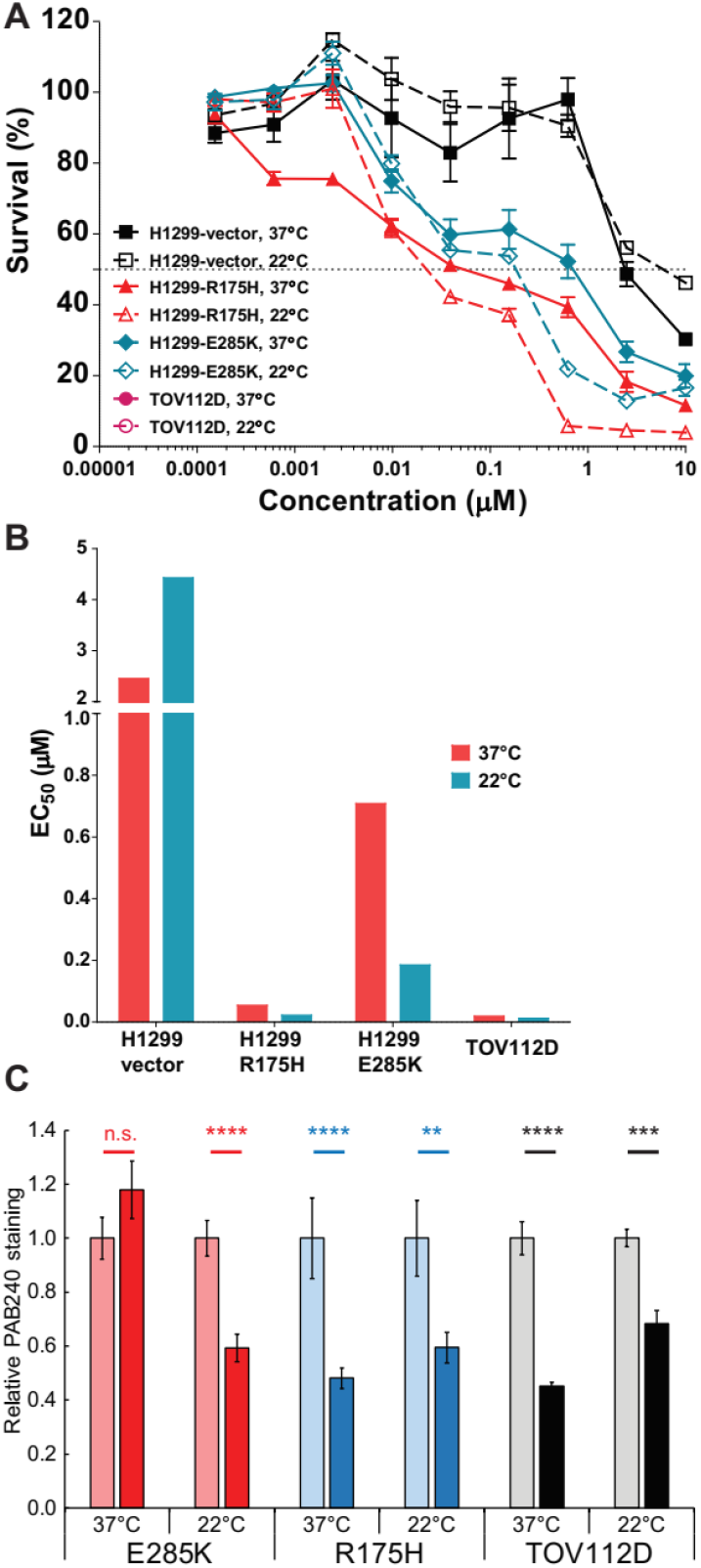
p53 folding landscape at lower temperature in cells. **(A)** Decreasing T synergizes with ZMC1 to kill cells expressing mutant p53. Cells were treated with 1 μM ZMC1, incubated at 37 °C or 22 °C for 4 h, incubated at 37 °C for 72 h, then assayed for viability by Calcein AM. **(B)** EC_50_ values from the curves in panel A. **(C)** Decreasing T synergizes with ZMC1 to refold mutant p53 in cancer cell lines. Experimental protocol is the same as in Fig. 4A. **, p < 0.01; ***, p < 0.001; ****, p < 0.0001; n.s., not significant.

### Restoration of p53 function in vivo

ZMCs are currently in pre-clincal development and represent a viable strategy to reactivate mutant p53 in the clinic. One of the attractive features of the ZMC program is that the spectrum of patients that will potentially respond to the drugs is known (those that harbor zinc-binding class p53 mutations). To determine if this spectrum should include individuals that have stability class mutations, we sought to obtain pre-clinical evidence for this using the xenograft tumor assay. We treated the animals with the tumors derived from the stable cell lines with V272M and E285K mutations with ZMC1 and found that the growth of the V272M tumors was inhibited but growth of E285K tumors was not (Fig. 6; SI Fig. 9), implying that ZMC1 can be used to reactivate certain p53 stability mutations, in addition to the zinc binding deficient mutations.

**Figure 6.**
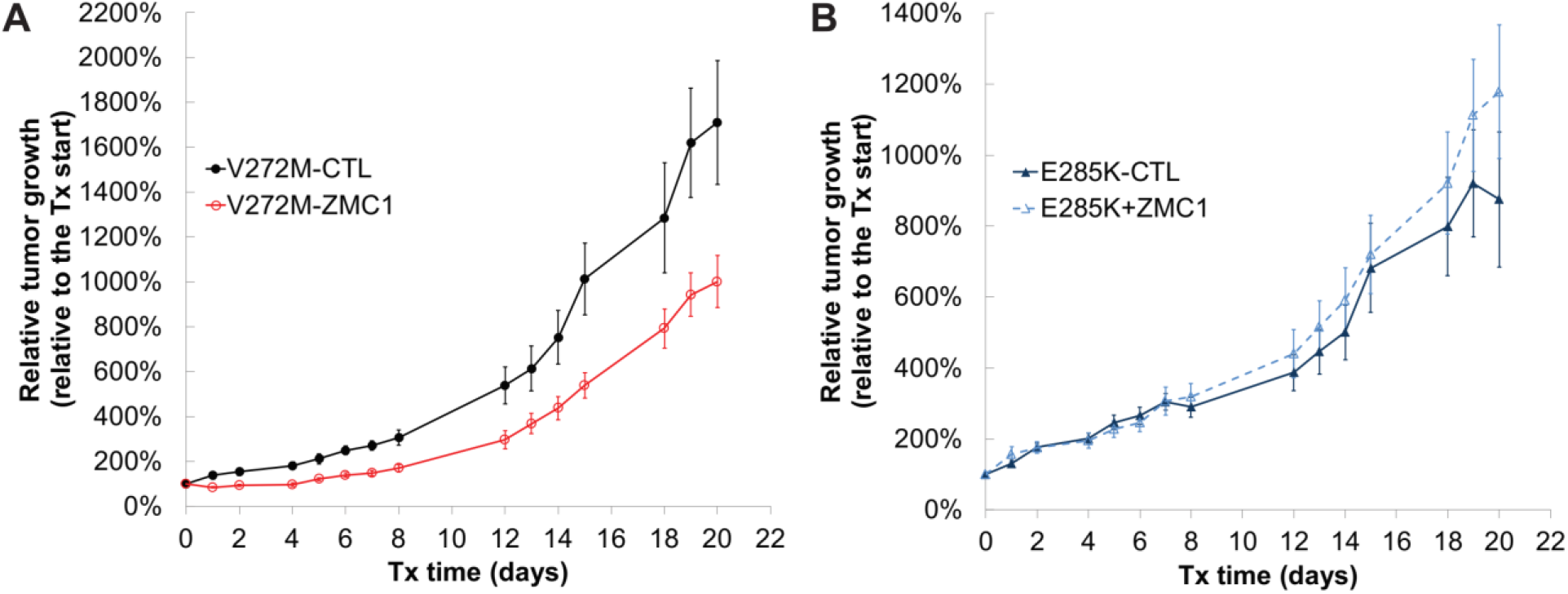
*In vivo* efficacy of ZMC1 in stability-class mutants V272M (A) and E285K (B). Mice bearing human xenograft tumors were treated with ZMC1 (5 mg/pk daily, IP) or DMSO vehicle.

### WT p53 is balanced between folded and unfolded states

We hypothesized from the energy landscape map that at physiological conditions of temperature and available zinc concentration, WT p53 may exist in approximately equal populations of folded and unfolded molecules. To address this question in context of the cell, we employed three cell lines expressing WT p53: MCF7, U2OS, and H460. We immunoprecipitated p53 protein from cell lysates with PAB240 or PAB1620 and blotted with the pan-p53 antibody DO-7 (Fig. 7). Because the two antibodies may have different affinities for p53, the percentages of unfolded and folded p53 cannot be determined quantitatively from this experiment. The intensities of the PAB240 bands, however, are comparable to those of the PAB1620 bands, suggesting that a significant fraction of WT p53 is unfolded in all three cell lines.

**Figure 7.**
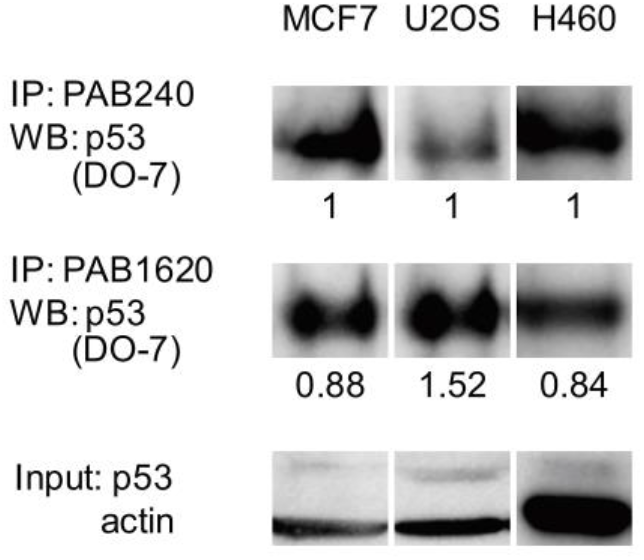
A significant fraction of WT p53 is unfolded in cells. WT p53 cell lines MCF7, U2OS, and H460 were lysed and immunoprecipitated using PAB240 or PAB1620 antibodies.

## Discussion

Our thermodynamic model of p53 folding is derived from two measurable properties: the free energy of DBD folding in the absence of metal, and the binding affinity of zinc to the folded protein. Of the 22 most common tumorigenic mutations examined here, all but three reduce stability by >1 kcal mol^−1^, decrease Zn^2+^ binding affinity by >10-fold, or both. The remaining three belong to the DNA contact class. These findings suggest that loss of thermodynamic stability and/or metal binding affinity play a dominant role in p53-related cancers, and underscore p53’s remarkable sensitivity to missense mutation at nearly every codon position.

In cells, ZMC1 activated three of the zinc-binding mutants (R175H, C176S, L194F) for cell killing (Table 2), PAB240-monitored refolding (Fig. 4A), and *PUMA*/*NOXA* transcription (Fig. 4C). Together with the zinc coordinating mutants that we previously characterized (C238S and C242S ^14^), five of the seven zinc-binding class tested responded to ZMC1 treatment, with P152L and R282Q being the exceptions. ZMC1 also failed to functionally rescue the mixed zinc-binding/stability mutant M237I. The energetic modeling does not consider structural effects, so it is expected that some mutants will remain inactive in the cell even if proper stability and zinc binding are restored by metallochaperones. For example, the R282Q mutation, being in the same helix as the DNA contact mutation R280K, is likely to perturb the structure of the DNA binding pocket, and ZMC1 appears to refold M237I (Fig. 4A) but not induce *PUMA*/*NOXA* transcription (Fig. 4C).

One of the primary predictions of the model is that increasing zinc concentration will refold stability class as well as zinc-binding class mutants. The archetypical example of small molecule-induced p53 stabilization is binding of PhiKan and similar compounds to the surface cavity on Y220C left by the Tyr220→Cys alteration ^28–31^. These drugs offer the potential of bio-orthogonality but bind to only a single p53 mutant and do so with (thus far) weak affinity (K_d_ = 1-100 μM). Zinc-induced stabilization, by contrast, takes advantage of an existing high-affinity site on p53 (K_Zn_ ~10^−15^ M) and can in principle refold any stability-class mutant, but suffers from lack of bio-orthogonality. As evidence for the potential breadth of ZMC therapy, treating cells with ZMC1 reactivated three stability-class mutants in cells (Y234A, V272M, and G245S ^14^) and in vivo (V272M), failed to reactivate two (Y234C and E285K). Y234C, like all the Tyr→Cys mutants that we tested, is refractory to zinc-induced stabilization most likely because of metal misligation in unfolded or partially folded states. E285K is the most unstable variant that we have characterized, which may explain why elevated zinc alone was insufficient for refolding without the additional stabilizing factor of reduced temperature. Nonetheless, the discovery that ZMC’s can reactivate mutants beyond just the class of zinc deficient mutation is a significant finding.

Previous work by our laboratory as well as that of Fersht established that the stability of DBD is low at 37 °C. The new insight offered by the current study is that DBD achieves this instability by a unique mechanism. In the absence of zinc, we find that WT apoDBD is much more unfolded than previously thought at body temperature (ΔG_apo_ = 6.7 kcal mol^−1^). This is not due to DBD being intrinsically disordered—it is quite stable at 10 °C—but to the anomalously high dependence of ΔG_apo_ on temperature. Offsetting DBD’s inherent propensity to unfold is its extraordinary affinity for zinc (K_Zn_ = 1.6 × 10^−15^ M), one of the highest yet reported for any eukaryotic protein. These biophysical data suggest that, in the absence of other cellular considerations such as chaperones and p53 binding partners, a dominant factor determining whether p53 is folded or unfolded is the available concentration of cytosolic zinc. At typical intracellular concentrations of available Zn^2+^ (10^−10^ M), our modeling predicts that the folded and unfolded populations are comparable—a prediction supported by conformation-specific antibody experiments (Fig. 7). This balance may explain why so many different missense mutations at nearly every codon position in the DBD gene are associated with loss of p53 function and cancer ^3^. More often than not, an amino acid substitution at any given position will decrease folding free energy rather than increase it, and loss of thermodynamic stability is the major cause of diseases that are caused by missense mutation of a single protein ^32^. To emphasize this point, nearly all mutants examined in this study destabilize DBD and/or decrease its zinc affinity. The remainder are DNA contact mutants, which can be reliably deduced from inspecting the X-ray crystal structure of DBD.

Why, then, might p53 have evolved with this unusual and precarious combination of high instability and zinc affinity? One explanation is that it constitutes a built-in failsafe to help rein in p53’s powerful cytotoxic activities should cellular checkpoint pathways become compromised. Carrying this scenario further, one might also speculate that it is a mechanism by which the cell can reversibly regulate p53 function by modulating its conformation through available zinc levels. Evidence for conformational regulation of p53 has existed for some time. Milner and Watson reported that fresh medium induced cell cycle and conformational changes in p53 ^33^. Hainaut, Milner, and colleagues reported that incubating cells and cell lysates with metal chelators can starve p53 of Zn^2+^ and induce the PAB240-binding form of p53, which can be rescued by re-introducing zinc ^13,34^. They also reported that oxidative agents when applied to cells can also induce the PAB240 conformation in WT p53. Oxidative agents increase intracellular free zinc levels by oxidizing the zinc binding cysteine residues on cytosolic metallothionein proteins, decreasing their affinity for zinc ^35^.

Approximately 10% of the proteins encoded by the human genome bind zinc; however, not until the last decade have researchers discovered that cytosolic zinc levels are in the picomolar range while the total cellular zinc is in the hundreds of micromolar ^24^ and more importantly, that many proteins are functionally regulated by zinc ^35^. This explains why there exists a complex repertoire of cellular homestatic genes consisting of cellular importers (ZIPs), antiporters (ZnTs) and cytosolic zinc buffers (metallothioneins). Proteins such tyrosine phosphatases are regulated by zinc, and these proteins typically have zinc affinities in the range of cytosolic free zinc ^36^. Although K_Zn_ of apoDBD is extremely low (1.6 × 10^−15^ M), the effective K_Zn_ at physiological temperature (K_Zn,eff_) is orders of magnitude higher owing to the inherent instability of apoDBD at 37 °C. K_Zn,eff_ is approximated by the product of K_Zn_ and K_apo_ (Eq. 4)—a value that our modeling suggests is close to cytosolic [Zn^2+^]_free_ (Fig. 3A). This suggests that physiologic perturbations in cytosolic zinc levels (100’s of pM), could in theory modulate zinc binding to p53 and hence its function. Moreover, given our findings that both apo and holo forms of WT p53 can be detected in cells, this suggests that p53 could potentially be regulated conformationally by zinc.

In conclusion, we have quantified the folding free energies and zinc binding affinities of the 20 most prevalent p53 mutations in cancer, many of which have not previously been characterized. Our thermodynamic modeling places the mutations into three distinct classes that will be useful to strafity patients for potential zinc metallochaperone treatment. We have demonstrated that ZMC1 treatment rescues the function of not only zinc binding class mutants (e.g. R175, C176, H179, C242), but also that of some stability class mutants (e.g. G245 and V272). Mutations in these six positions alone define a pool of 120,500 new cancer patients each year in the United States ^37^.

## Acknowledgements

We thank Dr. Arnold Levine for the p53^A138V^ cell line and Dr. Carol Prives for the luciferase reporter construct p21short. This work was supported by NIH grants R01 CA200800 and K08 CA172676 and from the Breast Cancer Research Foundation (to D.R.C).

## Materials and Methods

A detailed description of all materials, equipment, and methods in this study can be found in the SI Appendix.

### Protein purification and preparation of zinc solutions

DBDs (p53 residues 94 – 312) were expressed in *E. coli* and purified as previously described, and were >98% pure as judged by reducing SDS-PAGE stained with coomassie brilliant blue (11, 21). Apo proteins were generated as previously described (9). Full-length p53 was expressed as a fusion construct with a cleavable N-terminal Histag-ribose binding protein derived from *Thermoanaerobacter tengcongensis.* FL-p53 was purified by nickel-NTA chromatography (Qiagen) following manufacturers protocols. The ribose binding protein purification tag was then removed using human rhinovirus 3C protease, at which point FL-p53 was purified identically to DBD with the addition of a final clean-up step on a Supderdex S200 size exclusion column.

All *in vitro* experiments were performed in 50 mM Tris (pH 7.2), 0.1 M NaCl, 10 mM β-mercaptoethanol unless otherwise noted. ZnCl_2_ stocks were dissolved in 30 mM HCl and their concentrations determined by titration with 4-(2-pyridylazo)resorcinol using ɛ_500_ = 66,000 M^−1^ cm^−1^ for the PAR-Zn^2+^ complex. [Zn^2+^]_free_ concentrations were fixed at the indicated values by mixing 2 mM chelator (EDTA, N-(2-Hydroxyethyl)EDTA, EGTA, or diethylenetriaminepentaacetic acid) with 0.005 – 1.75 mM ZnCl_2_. [Zn^2+^]_free_ values were calculated using the MAXCHELATOR software suite (http://somapp.ucdmc.ucdavis.edu/pharmacology/bers/maxchelator/).

### Protein stability, zinc binding, and DNA binding assays

Urea denaturation experiments were performed as described ^12^. For fitting to the Gibbs-Helmholtz equation, samples were equilibrated at the indicated temperatures for 24 h before data collection, and curves were fit with a single linked *m*-value and all other parameters allowed to float. K_Zn_ competition assays were carried out by incubating unfolded apoDBD (in 6 M urea) at the indicated concentrations with 30 nM ZnCl_2_ and 30 nM FZ3 for 60 m at room temperature. FZ3 fluorescence was then scanned and K_Zn_ was obtained by fitting the data as described in SI Appendix. K_DNA_ values were obtained using 5’-Cy3 labeled oligonucleotides. Annealed oligonucleotides were mixed with 5 – 10,000 nM DBD on ice for 60 m. Fluorescence anisotropy was then measured and K_DNA_ was obtained by fitting the data as described in SI Appendix.

### Cell growth inhibition assays

Cell growth inhibition assay was performed by Calcein AM assay (Trevigen, MD). Variable temperature incubations were performed as described in the text. Statistical significance of the data, obtained from triplicate, independent samples, was calculated with Student’s *t*-test.

### Gene expression quantification

RNA was extracted using the RNeasy kit (Qiagen) and gene expressiong measured by quantitative RT-PCR, using the TaqMan gene expression assay system (Life Technologies).

### Immunofluorescent staining and immunoprecipitation

IF staining was performed as described previously ^26^ with additional details in Supplementary Information.

### Mouse experiments

Mice were housed and treated according to guidelines established by the Institutional Animal Care and Use Committee of Rutgers University, who also approved all mouse experiments. Nude mice (NCR nu/nu) were purchased from Taconic. Xenograft tumors were generated from the stable tumor cell lines H1299-V272M and H1299-E285K, (1 × 10^7^ cells/tumor site/mouse). Tumor dimensions were measured every 1-4 d and their volumes were calculated by using the formula: V = length × width^2^ × π / 6. Tumors (14 per group) were allowed to grow to 50 mm^3^ prior to daily administration of ZMC1 by intraperitoneal injection at the doses detailed in the text.

