## Supplemental Figures, Tables, and Methods for "Zinc shapes the folding landscape of p53 and establishes a new pathway for reactivating structurally diverse p53 mutants"

**Table of contents**

SI Fig. 1. Physical analysis validation data

SI Fig. 2. Free energy of folding versus buffered $[Zn^{2+}]_{free}$ for p53 DBD mutants

SI Fig. 3. DNA-binding of p53 DBD mutants

SI Fig. 4. Cell growth inhibition of p53 mutants by ZMC1

SI Fig. 5. Response of the p53 mutant cells to ZMC1.

SI Fig. 6. Refolding of p53 mutants after ZMC1treatement

SI Fig. 7. Binding of p53M237I to p21 promoter DNA

SI Fig. 8. Folding of p53 protein in V138 cells at 37°C with ZMC1

SI Fig. 9. Individual tumor growth curves for the processed data presented in Fig 6.

SI Table 1. DNA sequences used in protein-DNA binding experiments

SI Table 2. Oligonucleotides used to generate p53 mutants by site-directed mutagenesis

Extended Methods

References

Supplemental Figure 1

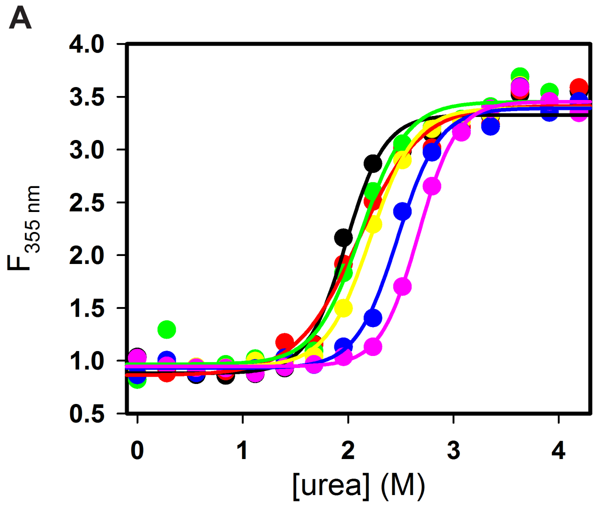


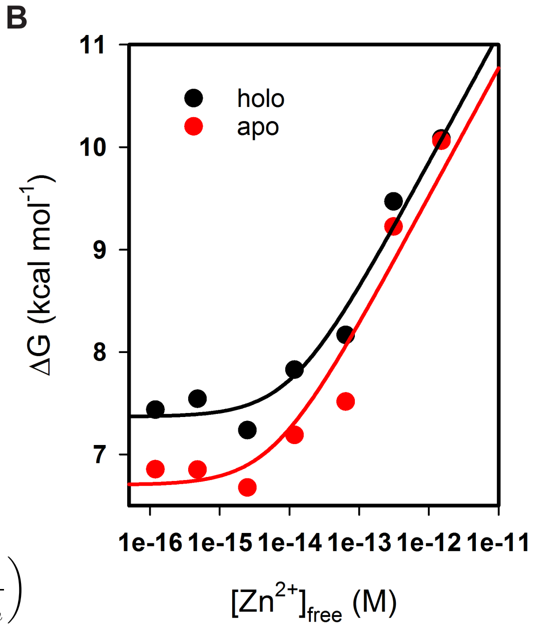


C

$$\Delta G_u(T) = \Delta H\left(1 - \frac{T}{T_m}\right) - \Delta C_p\left(T_m - T + T \cdot \ln\frac{T}{T_m}\right)$$

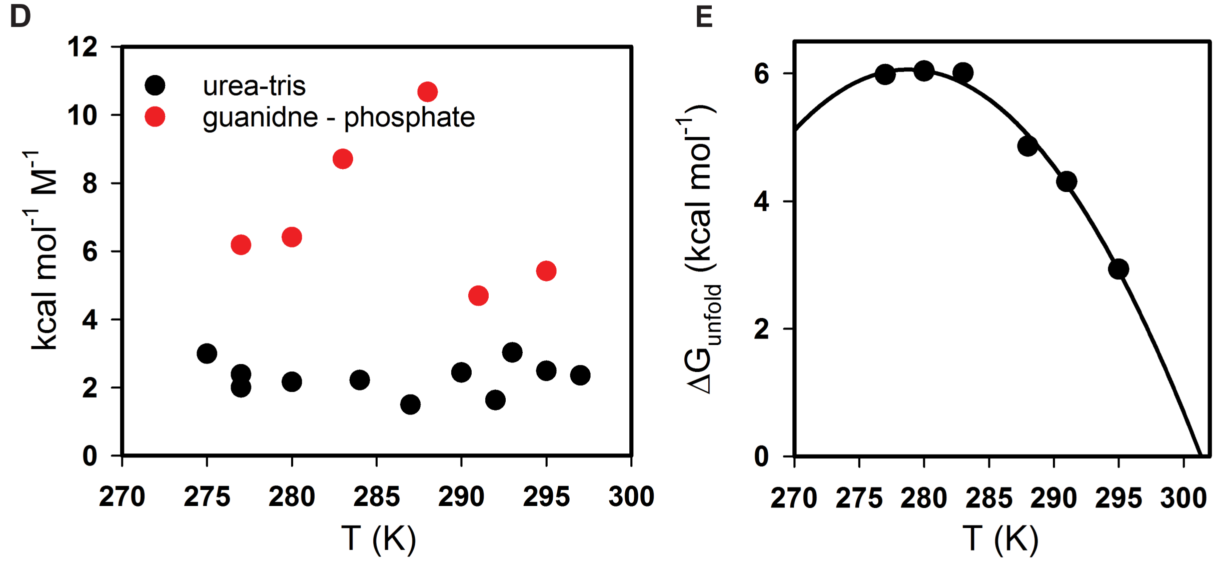


**Supplemental Figure 1. Physical analysis validation data. (A)** Urea melts of wt DBD as a function of buffered $[Zn^{2+}]_{free}$: $10^{-12}$ M (purple), $10^{-13}$ M (blue), $10^{-14}$M (yellow), $10^{-15}$ M (green), $10^{-16}$ M (red), 0 M (black). The m-values in kcal mol$^{-1}$ M$^{-1}$ of these fits are, in the same order: 3.1, 2.8, 2.6, 2.6, 1.9, and 3.0. **(B)** The relationship between ΔG and $[Zn^{2+}]_{free}$ is similar whether or not the protein is apoized. **(C)** The Gibbs-Helmholtz equation relates the Gibbs free energy change of a process to the enthalpy change (ΔH), the temperature (T), and the change in heat capacity ($\Delta C_p$). **(D)** m-values for individual urea and guanidine melts as a function of temperature do not indicate two-state behaviour. **(E)** Temperature dependence of apoDBD folding free energy, measured using guanidine denaturation and fit to the Gibbs-Helmholtz equation yields $\Delta H_m$ = 160 ± 12 kcal mol$^{-1}$, $T_m$ = 301 ± 1 K, and $\Delta C_p$ = 6.8 ± 1.1 kcal mol$^{-1}$ K$^{-1}$.

Supplemental Figure 2

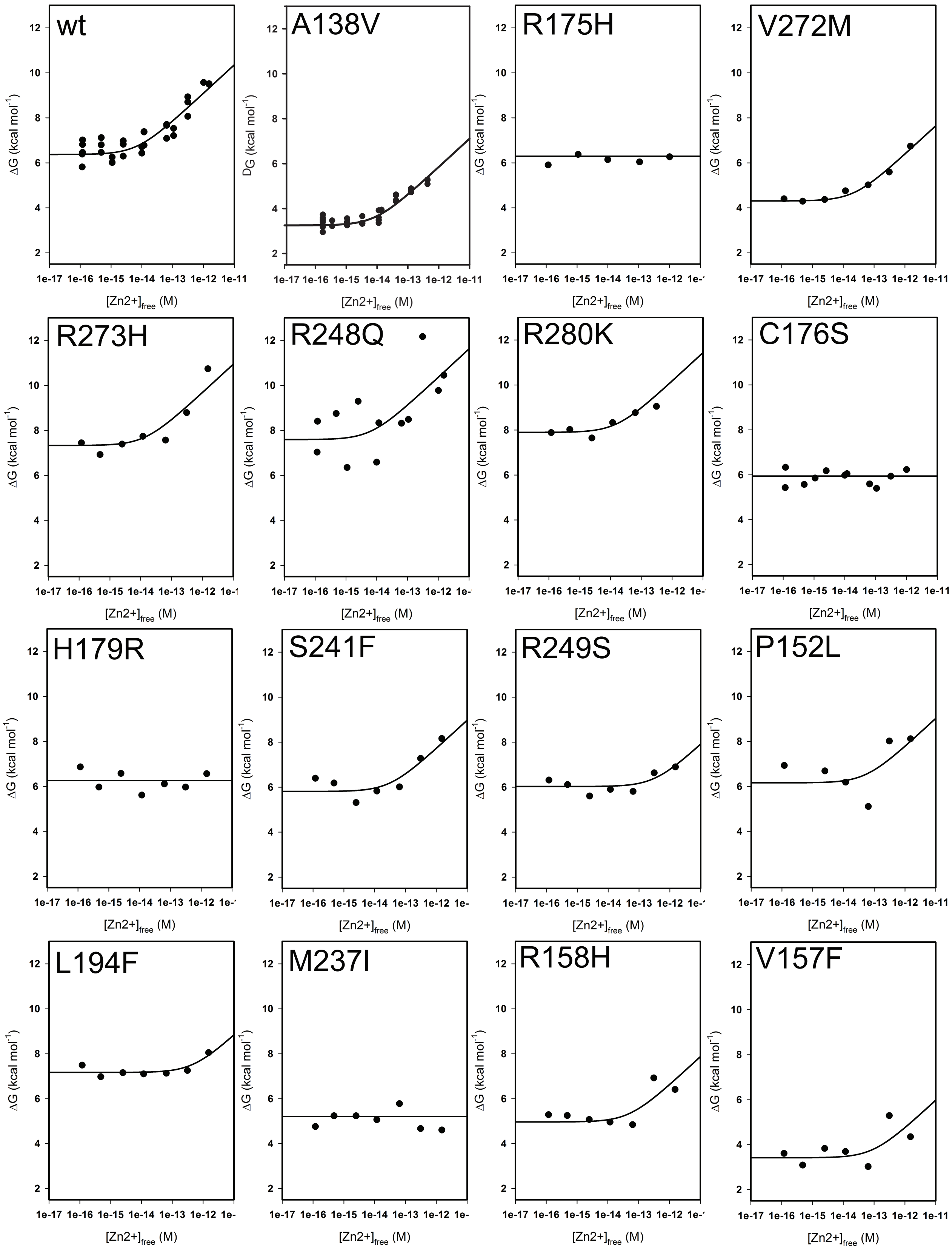

Supplemental Figure 2 - continued

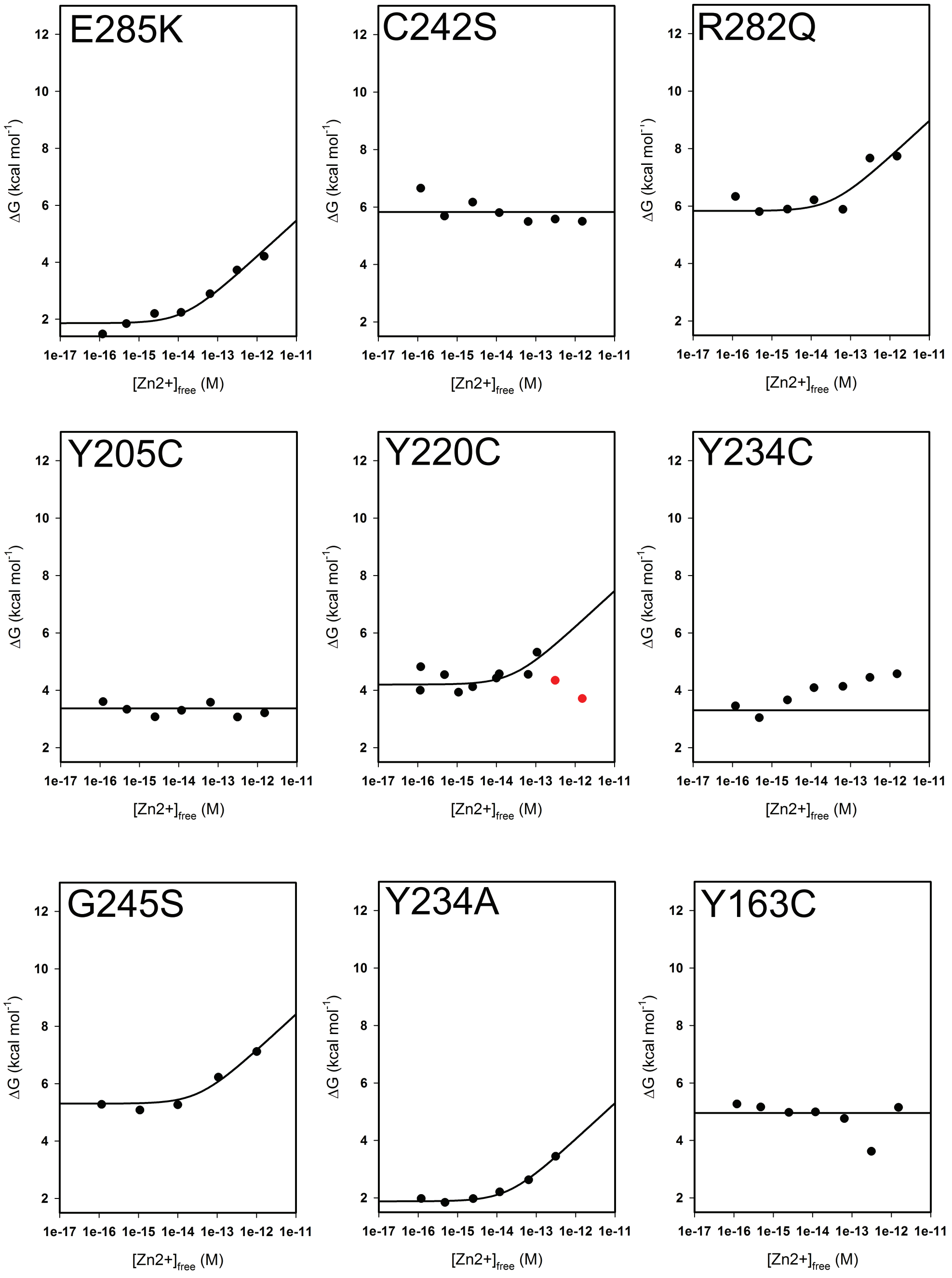


**Supplemental Figure 2. Free energy of folding versus buffered $[Zn^{2+}]_{free}$ for p53 DBD mutants.** Folding energies were determined from guanidine melts of DBD mutants in EDTA-buffered $[Zn^{2+}]_{free}$, and were used to determine the zinc-free folding energies and zinc affinities presented in Figure 3 C and Table 1. Red points indicate outliers excluded from analysis.

Supplemental Figure 3

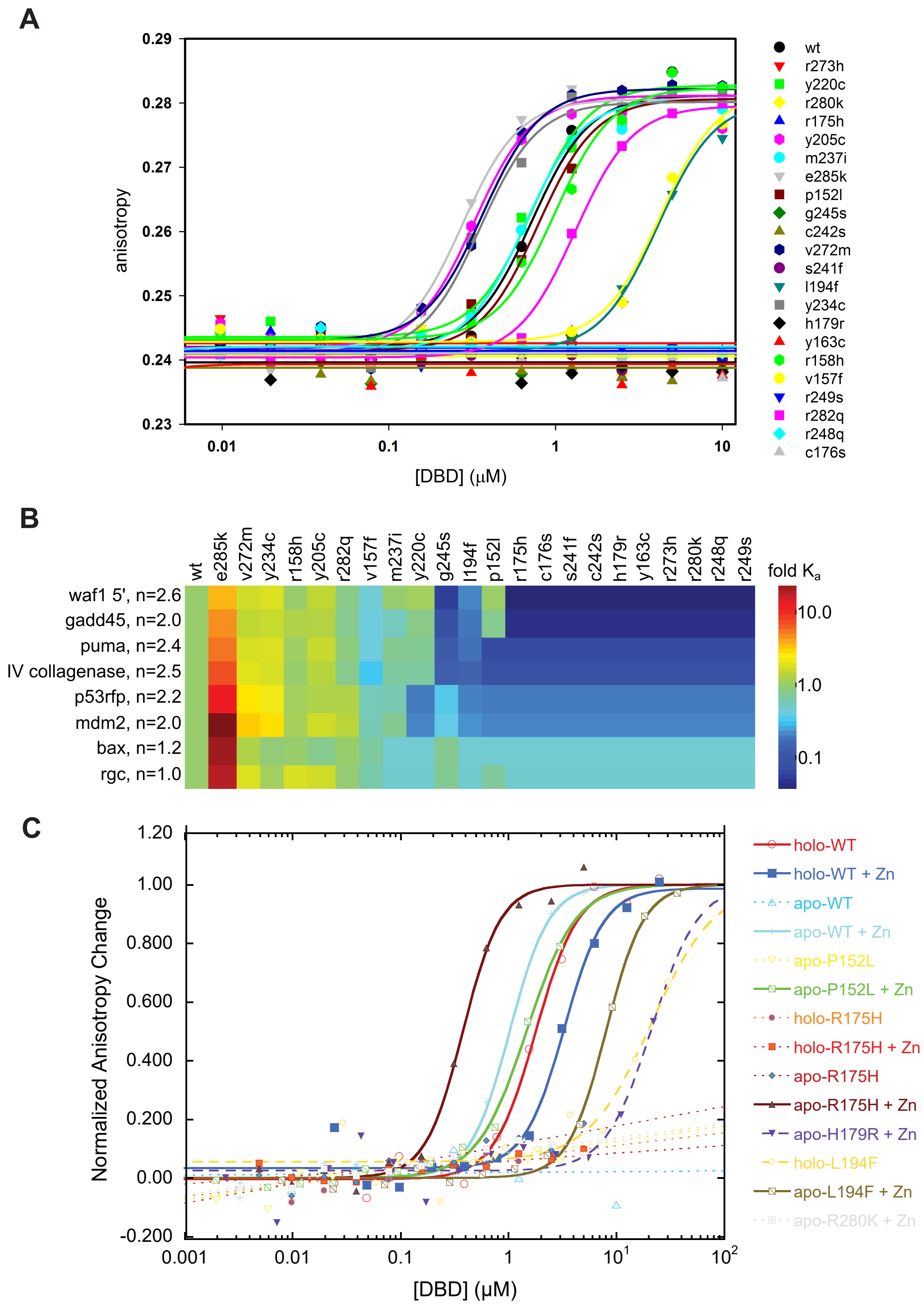


**Supplemental Figure 3. DNA-binding of p53 DBD mutants. (A)** Representative fits for un-apoized DBD mutants binding Cy3-labeled p53RE oligonucleotides. **(B)** Heatmap of mutant DBD affinity for DNA oligos relative to wt-DBD. Oligonucleotide names and Hill-parameter on the left. Sequences in Table S1. **(C)** Representative curves illustrating restoration of DNA-binding (waf1) for DBD mutants by EGTA-buffered $[Zn^{2+}]_{free}$.

Supplemental Figure 4

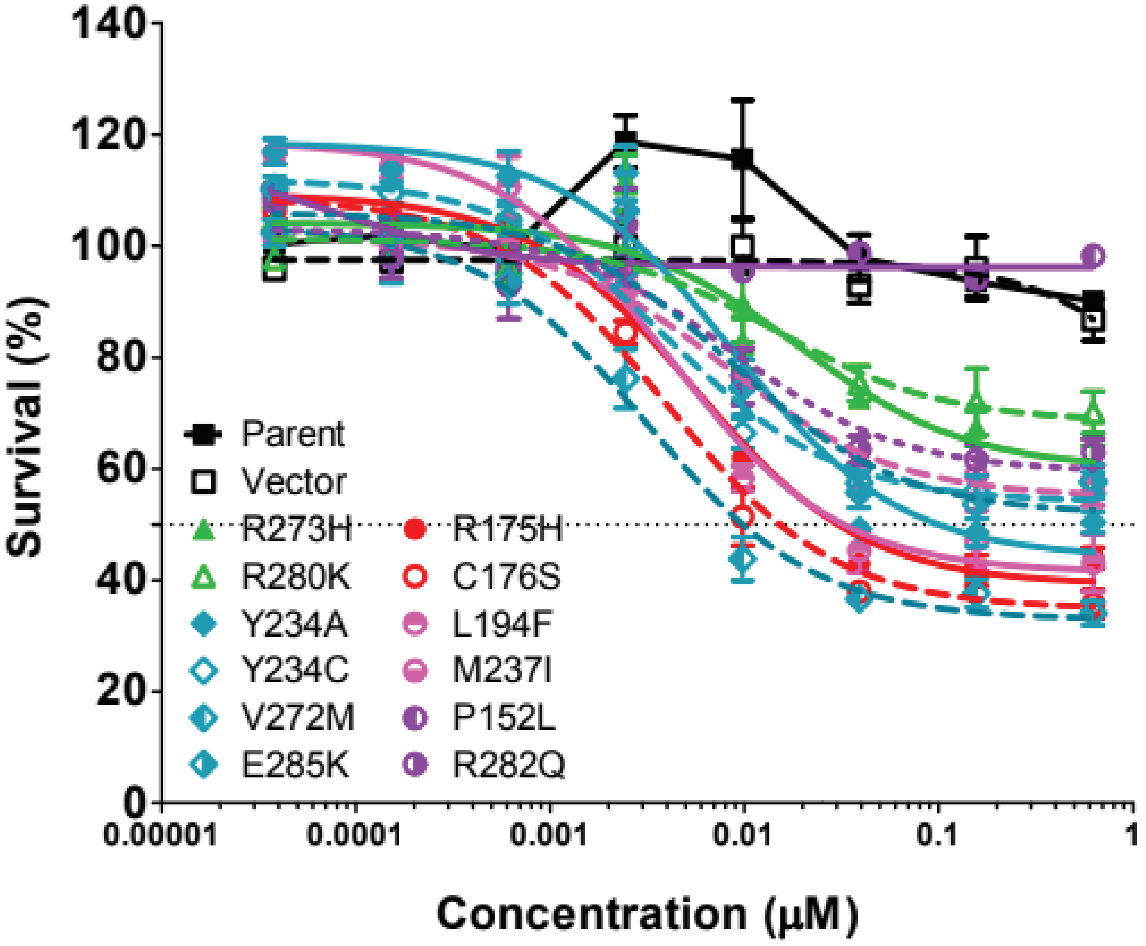


**Supplemental Figure 4. Cell growth inhibition of p53 mutants by ZMC1.** 12 common p53 mutants were generated by site-directed mutagenesis and expressed in p53-null H1299 cells. These were treated with ZMC1 and the cell growth inhibition was measured by Calcein AM assay. The $EC_{50}$ values were calculated using non-fit curves by GraphPad Prism software and summarized in Table 2.

Supplemental Figure 5

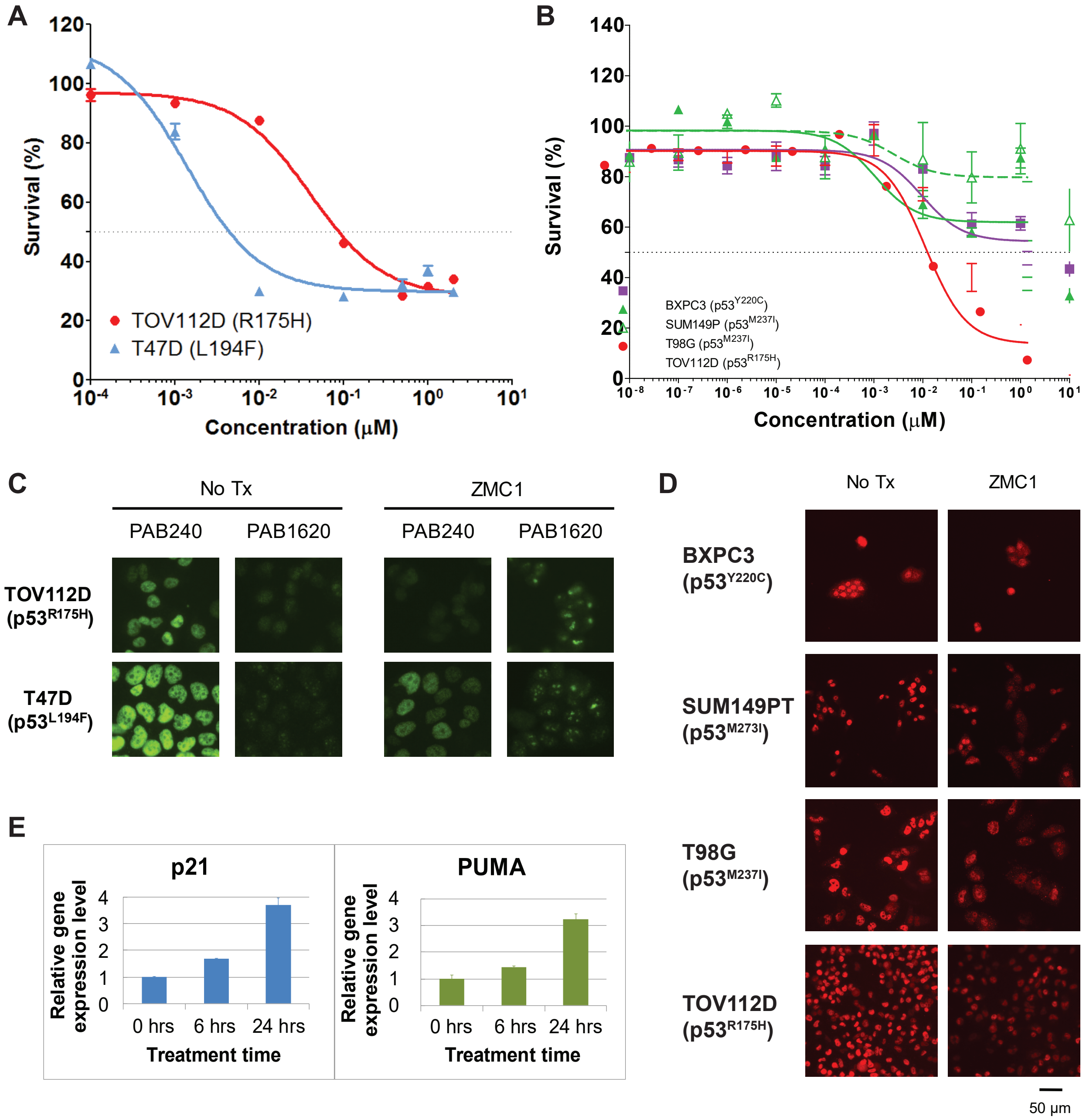


**Supplemental Figure 5. Response of the p53 mutant cells to ZMC1. (A)** T47D (p53L194F) cells are sensitive to ZMC1 treatment in a cell viability assay. **(B)** BXPC3 (p53Y220C), SUM149P (p53M237I) and T98G (p53M237I) cells are not sensitive to ZMC1 in a cell viability assay. **(C)** The p53 mutant L194F was refolded after ZMC1 treatment. The cells were treated with 1 μM ZMC1 for 6 hours, followed by IF using p53 antibodies PAB240, which binds the unfolded conformation, and PAB1620, which binds the native conformation. **(D)** p53 mutant Y220C was not refolded by ZMC1, but M237I was (IF with PAB240). **(E)** The gene expression of the p53 target genes *p21* and *PUMA* was increased after ZMC1 treatment in cells expressing p53 L194F.

Supplemental Figure 6

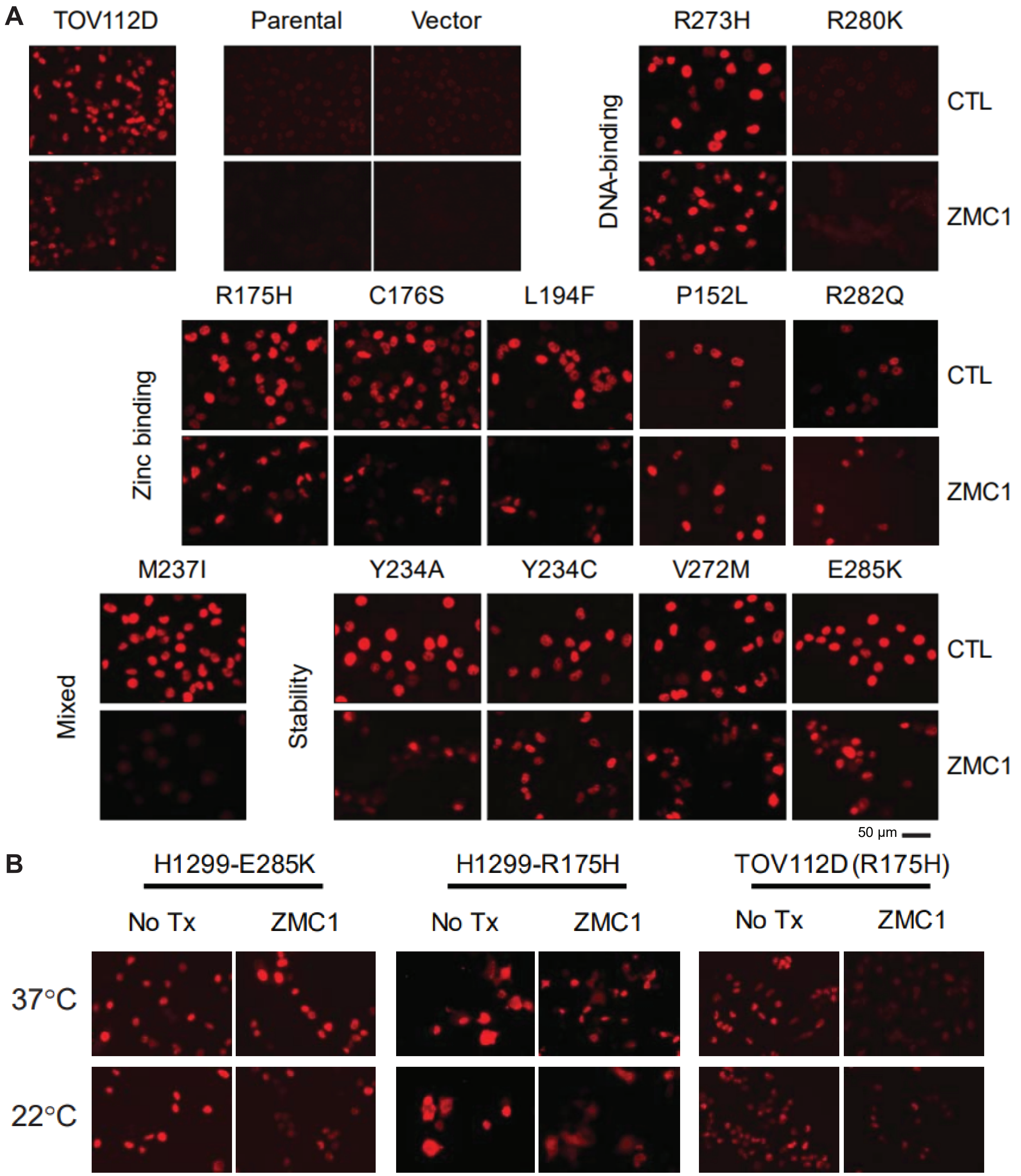


**Supplemental Figure 6. Refolding of p53 mutants after ZMC1treatement. (A)** H1299 cells were stably transfected with 1 of 12 p53 mutants or an empty vector, and treated with ZMC1 for 4 hours. p53 conformation was probed by IF using PAB240, which binds selectively to unfolded p53. TOV112D (p53 R175H) and the parental cell line were used as unfolded and native conformation controls respectively. **(B)** p53 refolding of R175H and E285K with ZMC1 and different incubation temperatures

Supplemental Figure 7

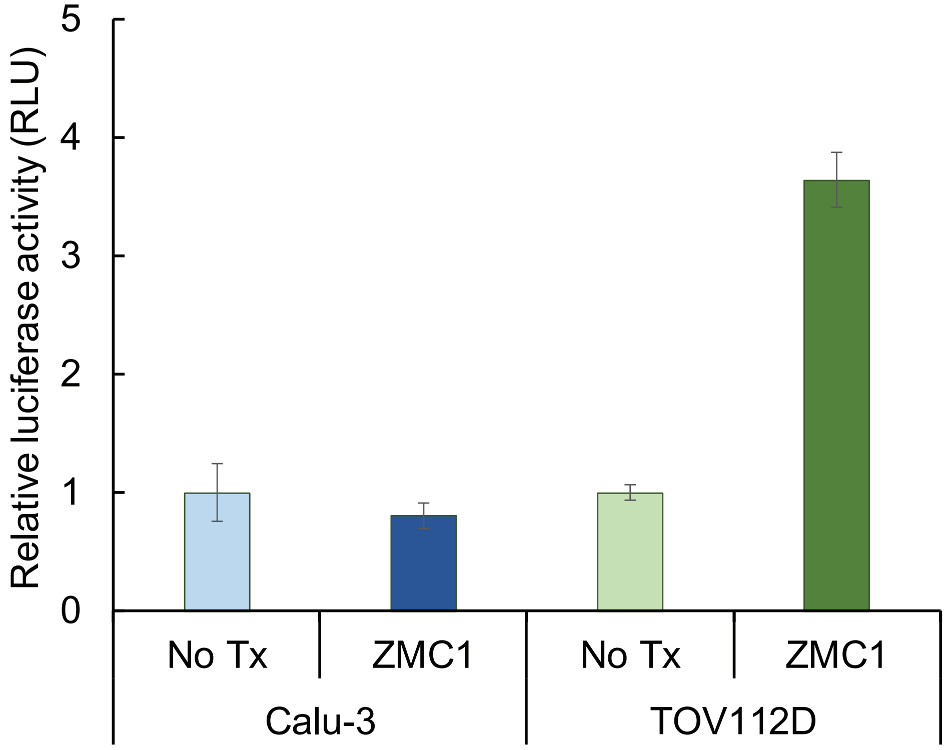


**Supplemental Figure 7. Binding of p53M237I to p21 promoter DNA.** Calu3 (p53M237I) cells were transfected with the p53RE from the p21 promoter sequence (p21short) or a negative control (Basic). The cells were treated with ZMC1 for 48 hours, then assayed for luciferase activity. TOV112D (p53R175H) is used as a positive control.

Supplemental Figure 8

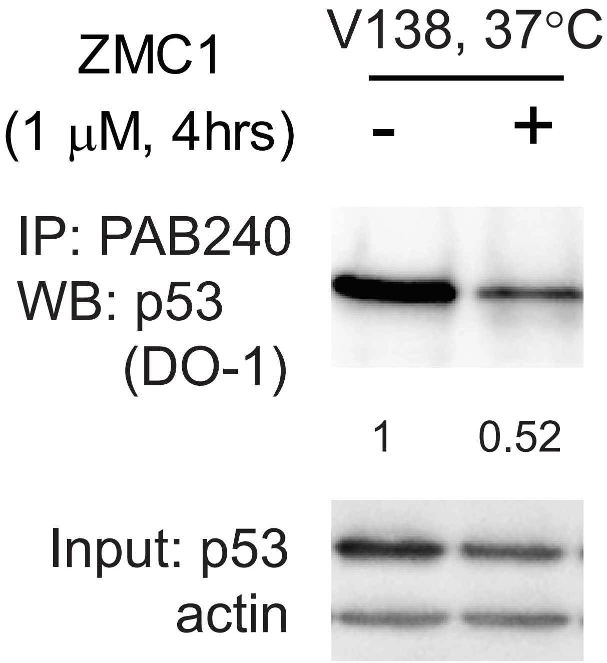


**Supplemental Figure 8. Folding of p53 protein in V138 cells at 37°C with ZMC1.** Cells were treated with 1 μM for 4 hours, and protein conformation was determined by IP of the unfolded protein.

Supplemental Figure 9

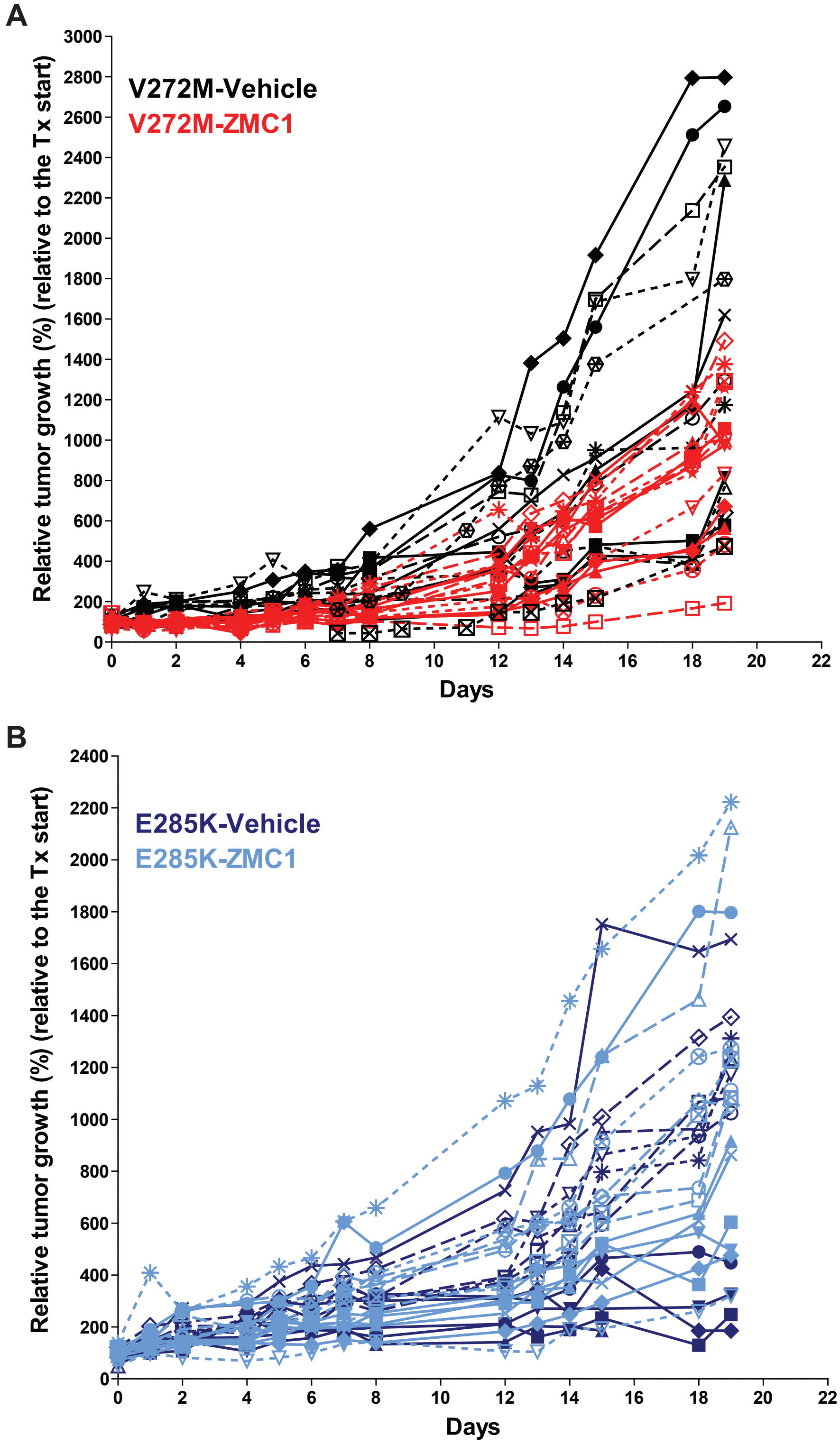


**Supplemental Figure 9. Individual tumor growth curves for the processed data presented in Fig 6.** Tumor growth shows the in vivo efficacy of ZMC1 in stability-class mutant V272M **(A)** but not in E285K **(B)**.

| **Supplemental Table 1: DNA sequences used in protein-DNA binding experiments** | | |
|---|---|---|
| **Name** | **Sequence (5'-3')** | **5' Cy3 label** |
| gadd45 (forward) | gaacatgtctaagcatgctg | + |
| gadd45 (complement) | cagcatgcttagacatgttc | - |
| puma (forward) | ctgcaagtcctgacttgtcc | + |
| puma (complement) | ggacaagtcaggacttgcag | - |
| mdm2 (forward) | ggtcaagttcagacacgtcc | + |
| mdm2 (complement) | ggacgtgtctgaacttgacc | - |
| rgc (forward) | ggacttgcctggccttgcct | + |
| rgc (complement) | aggcaaggccaggcaagtcc | - |
| p53rfp (forward) | agacaggtcctgacaagcag | + |
| p53rfp (complement) | ctgcttgtcaggacctgtct | - |
| waf1 5' (forward) | gaacatgtcccaacatgttg | + |
| waf1 5' (complement) | caacatgttgggacatgttc | - |
| waf1 3' (forward) | gaagaagactgggcatgtct | + |
| waf1 3' (complement) | agacatgcccagtcttcttc | - |
| bax (forward) | agacaagcctgggcgtgggc | + |
| bax (complement) | gcccacgcccaggcttgtct | - |
| IV collagenase (forward) | agacaagcctgaacttgtct | + |
| IV collagenase (complement) | agacaagttcaggcttgtct | - |
| egfr (forward) | gagctagacgtccgggcagcccc | + |
| egfr (complement) | ggggctgcccggacgtctagctc | - |

| Supplemental Table 2: Oligonucleotides used to generate p53 mutants by site-directed mutagenesis | | |
|---|---|---|
| **p53 mutation** | **Forward primer (5′–3′)** | **Reverse primer (5′–3′)** |
| Vector | purchased | purchased |
| **Zinc-binding mutants** | | |
| R175H | gttgtgaggcactgcccccac | ctccgtcatgtgctgtgac |
| C176S | gtgaggcgctccccccaccat | aacctccgtcatgtgctgtgac |
| L194F | tcctcagcattttatccgagtggaag | ggggccagaccatcgcta |
| P152L | tccacacccctgcccggcacc | atcaacccacagctgcacagggc |
| R282Q | gggagagaccagcgcacagag | aggacaggcacaaacacg |
| **Stability mutants** | | |
| Y234A | caccatccacgccaactacatgtgtaac | gtacagtcagagccaacc |
| Y234C | accatccactgcaactacatg | ggtacagtcagagccaac |
| V272M | cagctttgagatgcgtgtttg | ttccgtcccagtagattac |
| E285K | ccggcgcacaaaggaagagaa | tctctcccaggacaggcac |
| **Mixed mutant** | | |
| M237I | acaactacatctgtaacagttcct g | agtggatggtggtacagt |
| **DNA-binding mutants** | | |
| R273H | purchased | purchased |
| R280K | tgtcctgggaaagaccggcgc | ggcacaaacacgcacctcaaagc |

### Extended Methods

***Reagents.*** FluoZin-3, tetrapotassium salt (FZ3) and cell culture media were purchased from Life Technologies Corporation (Norwalk, CT). Cell lines were purchased from American Type Culture Collection (Manassas, VA). Cy3-labeled (PAGE purified) and unlabeled (HPLC purified) p53 recognition element oligonucleotides were purchased from Eurofins Genomics (Louisville, KY). All other chemicals were purchased from Sigma-Aldrich (St. Louis, MO) and were >98% pure or better.

***Protein expression and purification.*** Recombinant p53 DNA-binding domains were expressed and purified as previously described and were >98% pure as judged by reducing SDS-PAGE stained with Coomassie brilliant blue [1,2]. Apo proteins were generated as previously described [1]. Full length p53 (FL-p53) was expressed as a fusion construct along with an N-terminal 6xHis tag-ribose binding protein derived from *Thermoanaerobacter tengcongensis* and human rhinovirus 3C protease site in pCDFDuet-1 vector (EMD Millipore, Darmstadt, Germany). Chemically competent BL21(DE3) cells were transformed with the expression plasmids, plated on lysogeny broth (LB) agar plates containing 50 μg/mL streptomycin, and grown at 37 ˚C overnight. Isolated colonies then picked and grown in LB + 50 μg/mL streptomycin at 37 ˚C with 200 RPM shaking until OD600 = 0.6. The temperature was then dropped to 18 ˚C and the cultures induced with 20 mg/L isopropyl-β-D-thiogalactoside overnight. Cells were harvested by centrifugation, resuspended in resuspension/wash buffer (20 mM Tris pH 7.2, 300 mM NaCl, 10 mM imidazole, and 10 mM β-mercaptoethanol), and lysed enzymatically. Insoluble material was pelleted, and the supernatant loaded on to a Ni-

NTA (Qiagen, Valencia GA) column pre-equilibrated with resuspension/wash buffer. After washing, the sample was eluted with 20 mM Tris pH 7.2, 300 mM NaCl, 250 mM imidazole, and 10 mM β-mercaptoethanol. Protein-containing fractions were pooled, dialyzed against 20 mM Tris, 150 mM NaCl, 10 mM β-mercaptoethanol, and the tags removed by incubation with GST-tagged HRV 3C protease (0.05-0.1 mg protease/mg p53) for ~18 hrs at 4 ˚C. The protein was further purified by heparin pseudoaffinity chromatography (0.15-1 M NaCl gradient) on a Heparin HiTrap column (GE Healthcare Life Sciences, Pittsburgh, PA). The final protein was >90% pure by SDS-PAGE stained with Coomassie brilliant blue. FL-p53 protein was folded by circular dichroism (CD) spectroscopy, exhibited the expected change in CD spectrum when stripped of $Zn^{2+}$, bound p53 consensus sequence DNA by electrophoretic mobility shift assay, eluted >90% as a monodisperse tetramer by gel filtration, and had identical $Zn^{2+}$-content per monomer relative to purified DBD. Proteins were flash frozen on dry ice and stored at -80 °C until use. *In vitro* experiments were conducted in 50 mM Tris pH 7.2, 100 mM NaCl, 10 mM β-ME at 10 ˚C unless otherwise noted.

***Preparation of $Zn^{2+}$-solutions.*** $ZnCl_2$ stocks were dissolved in 30 mM HCl and stored at room temperature. Concentration of $Zn^{2+}$ stock solutions were determined by incubation with 150 μM colorimetric $Zn^{2+}$ indicator 4-(2-pyridylazo)resorcinol (PAR) using $\epsilon_{500} = 6.6 \times 10^4$ for the PAR-$Zn^{2+}$ complex in buffer. FZ3 stocks were dissolved in $ddH_2O$ and stored at -20 °C until use. Concentration of FZ3 stocks were determined by equivalence point titration with $Zn^{2+}$ in buffer. Concentrations of $Zn^{2+}$-chelators (EDTA, HEDTA, EGTA, and DHPTA) were determined by titrating the chelator solutions against pre-formed PAR-$Zn^{2+}$ complex and measuring the decrease in absorbance in buffer. For intrinsic fluorescence measurements and urea melts, $[Zn^{2+}]_{free}$ was calculated using the program MaxChelator as previously described [3]. Chelator concentrations were always 2 mM, and $ZnCl_2$ concentrations ranged from 5-1750 μM, which was always in great excess of protein concentration (1 μM). Example calculation shown in Table S2.

***$Zn^{2+}$ $K_d$ measurements by intrinsic fluorescence.*** Apoized DBDs or FL-p53 were incubated with chelator/$ZnCl_2$ mixtures to maintain the indicated buffered $[Zn^{2+}]_{free}$ at 10 °C for 16 h. We observed a slightly increased fluorescence at 306 nm for DBD's when $Zn^{2+}$-bound, and a slightly increased fluorescence at 350 nm for FL-p53 when $Zn^{2+}$-bound. Measurements were taken on a Fluoromax-4 spectrofluorometer in a 5 mm x 5 mm quartz cuvette with $\lambda_{ex}$ = 280 nm.

Results for DBDs were fit to a single-site binding equation:

Eqn 1.
$$y = y0 + \frac{AL}{K_d + L}$$

where y is the measured fluorescence, y0 is the baseline value, A is the amplitude, L is the concentration of free ligand, and $K_d$ is the dissociation constant.

To determine if there was any cooperatively in the tetramer, that data was fit to a single-site Hill-binding equation:

Eqn 2.
$$y = y0 + \frac{AL^n}{K_d^n + L^n}$$

where y is the measured fluorescence, y0 is the baseline value, A is the amplitude, L is the concentration of free ligand, and $K_d$ is the mass-action dissociation constant, n is the Hill-parameter. Data were converted to fraction bound, and representative traces and fits shown for display. $K_d$ values and Hill-parameters were averaged from independent trials.

***$Zn^{2+}$ $K_d$ measurements by competition.*** Apo DBDs unfolded in 6 M urea at the indicated concentrations were incubated with 30 nM $ZnCl_2$ + 30 nM FZ3 at room temperature for 60 min in black polystyrene 96-well plates and the fluorescence measured using a SpectraMax i3x plate reader ($\lambda_{ex}$ = 500 nm, $\lambda_{em}$ = 540 nm) (Molecular Devices, LLC, Sunnyvale, CA). The data were plotted on a log axis and fit to a sigmoid to measure $IC_{50}$:

Eqn 3.

$$y = \frac{A}{1 + e^{-\left(\frac{x - x0}{b}\right)}}$$

Where y is the measured fluorescence, A is the curve amplitude, x is log[DBD], x0 is the $IC_{50}$, and b is an empirical steepness parameter. That $IC_{50}$ was then used in the Munson-Rodbard solution to the Cheng-Prussof equation to calculate $K_d$ [4]:

Eqn 4.

$$K_i = \frac{IC_{50}}{1 + \frac{L_T(y_0+2)}{2K_d(y_0+1)} + y_0} - K_d\frac{y_0}{y_0 + 2}$$

where $K_i$ is the dissociation constant of the DBD, $L_T$ is the total concentration of FZ3, $K_d$ is the dissociation constant of FZ3 for $Zn^{2+}$ (15 nM per manufacturer), and $y_0$ is the ratio of bound FZ3 to free FZ3 in the absence of DBD (15 nM/15 nM under these conditions).

***Urea Melts as a function of Zn²⁺.*** Buffer solutions containing either 0 or 5 M urea were mixed with a Hamilton Microlab 540B dispenser (Reno, NV) to make a linear gradient of urea concentrations. Buffers were then mixed with DBD and chelator/$ZnCl_2$ solutions to yield the indicated concentrations of urea, 1 μM DBD, 2 mM chelator, and 5-1750 μM $ZnCl_2$ to buffer the $[Zn^{2+}]_{free}$ to the indicated level as described above. Urea concentrations were determined by refractive index measurements [5]. Samples were incubated for 16 h, and fluorescence measured using either a Fluoromax-4 spectrofluorometer (Horiba Scientific, Edison, NJ) in a 5 mm x 5 mm quartz cuvette or in a SpectraMax i3x plate reader (Molecular Devices, LLC, Sunnyvale, CA) in 96-well UV-Microplates (Thermo Scientific, Waltham, MA) with comparable results ($\lambda_{ex}$ = 280 nm, $\lambda_{em}$ = 355 nm). The resultant curves were fit to the 2-state linear extrapolation model with linear baselines according to the equation [6,7]:

eqn 5.
$$F = \frac{y_U + s_U[D] + (y_N + s_N[D])e^{\frac{\Delta G_0 - m[D]}{RT}}}{1 + e^{\frac{\Delta G_0 - m[D]}{RT}}}$$

Where F is the measured fluorescence, $y_N$ is the y-intercept of the native baseline, $s_N$ is the slope of the native baseline, [D] is the concentration of denaturant, $y_U$ is the y-intercept of the unfolded baseline, $s_U$ is the slope of the unfolded baseline, $\Delta G_0$ is the folding energy in the absence of denaturant, m is the empirical m-value, R is the ideal gas constant, and T is the absolute temperature. To increase the accuracy of our measurements, we pooled m-values for all valid melts conducted for all naturally occurring mutants (3.10 ± 0.12 kcal $mol^{-1}$ $M^{-1}$, n = 213, mean ± SE) and calculated $\Delta G_0$ from the less error-prone $C_m$ value ($\Delta G_0$/m from each fit) using:

eqn 6.
$$\Delta G_0 = C_m \cdot m_{pooled}$$

Where $\Delta G_0$ is the folding energy in the absence of denaturant, $C_m$ is the concentration of denaturant at which the protein is 50% unfolded, and m is the pooled m-value.

Those measured ΔG values were then plotted as a function of $[Zn^{2+}]_{free}$, and fit to the substrate stabilization model [8]:

eqn 7.
$$\Delta G = \Delta G_{apo} + RTln(1 + \frac{[L]}{K_d})$$

Where ΔG is the measured folding energy, $\Delta G_{apo}$ is the folding energy in the absence of ligand, R is the ideal gas constant, T is the absolute temperature, [L] is the concentration of free ligand, and $K_d$ is the dissociation constant of the native protein for the ligand. All data for each mutant were pooled and a single global fit performed per mutant.

***Urea Melts as a function of T.*** Buffer solutions containing 2 mM EDTA, 1 μM apoDBD, and either 0 or 4 M urea were mixed as above and incubated at the indicated temperature overnight. Identical experiments were conducted in 50 mM phosphate buffer pH 7.2, 100 mM NaCl, 10 mM βME, 2 mM EDTA with 0-2 M guanidine hydrochloride. Melts were evaluated as above with eqn 5 and 6, using the pooled m-value for urea as above, and the pooled m-value for guanidine (7.01 ± 0.92 kcal $mol^{-1}$ $M^{-1}$, n = 6, mean ± SE). Measured $\Delta G_0$ were plotted as a function of temperature and fit to the Gibbs-Helmholtz equation:

eqn 8. $$\Delta G = \Delta H_m(1 - \frac{T}{T_m}) - \Delta C_p[T_m - T + Tln\frac{T}{T_m}]$$

Where ΔG is the free energy of unfolding at temperature T, $\Delta H_m$ is the enthalpy of unfolding at the $T_m$, $T_m$ is the temperature at which 50% of the protein is unfolded in the absence of denaturant, and $\Delta C_p$ is the change in constant pressure heat capacity from the folded to unfolded protein.

***DNA $K_d$ measurements.*** 5'-Cy3 labeled oligonucleotides with sequences of p53 recognition elements taken from the promoter regions of different p53 target genes were annealed to their unlabeled reverse complement by heating at 95 ˚C with a slight excess of unlabeled DNA for 2 min and slowly cooling to room temperature over 45 min (30 mM HEPES pH 7.5, 100 mM potassium acetate 100 μM/125 μM labeled/unlabeled). Annealed oligos (50 nM) were incubated with 5-10,000 nM DBD on ice for 1 h in 50 mM Tris pH 7.2, 100 mM NaCl, 1 mM TCEP, and 0.005% Tween-20 in black 96 well plates, and uncalibrated fluorescence anisotropy measured using SpectraMax i3x equipped with rhodamine fluorescence polarization module (G-factor = 1, $\lambda_{ex}\backslash\lambda_{em}$ = 535 nm/595 nm) (Molecular Devices, Sunnyvale, CA). Data for all trials of all mutants for a given sequence were subjected to global curve fitting with eqn 2, linking A and n across all trials. Sequences are given in Table S3.

***Reactivation of DNA binding in buffered $Zn^{2+}$.*** In EGTA-buffered $Zn^{2+}$ solutions were prepared as described above. 20,000-1 nM apo- or holo-DBD was incubated with 5 nM annealed 5'-Cy3 labeled oligonucleotides (described above) in 20 mM HEPES pH 7.2, 150 mM NaCl, 1 mM TCEP, 0.005% Tween-20, and EGTA-buffered $Zn^{2+}$ on ice for 1h in 96-well black plates. Fluorescence anisotropy was measured as described above, and the resulting curves were fit using eqn 2.

***Energy landscape plots.*** The zinc-free folding energy of DBD mutants as a function of temperature was calculated using the Gibbs-Helmholtz equation to determine temperature dependence (eqn 8), and the difference in zinc-free folding energy at a reference temperature to adjust for mutant stability.

eqn 9.

$$\Delta G_{apo,mut} = \Delta H_m\left(1-\frac{T}{T_m}\right) - \Delta C_p\left(T_m - T + Tln\left(\frac{T}{T_m}\right)\right) - \left(\Delta G_{ref,WT} - dG_{ref,mut}\right)$$

Where $\Delta G_{apo,mut}$ is the zinc-free folding energy of a given mutant at a given temperature, $\Delta H_m$ is the enthalpy of unfolding of wild-type DBD at $T_m$, $T_m$ is the temperature at which 50% of the wild-type DBD is unfolded in the absence of both denature and zinc, $\Delta C_p$ is the change in constant-pressure heat capacity from the folded to unfolded wild-type

DBD, and $\Delta G_{ref,WT}$ and $\Delta G_{ref,mut}$ are the free energy of unfolding of wild-type and mutant DBD under the same conditions.

The fraction of the mutant DBD that is folded in the absence of zinc as a function of temperature is therefore given by

Eqn 10 $$K_{apo,mut} = e^{\frac{\Delta G_{apo,mut}}{RT}}$$

Where R is the gas constant.

The folding landscape is subsequently calculated from the four-state model, assuming that the unfolded DBD only binds a single $Zn^{2+}$ ion.

Eqn 11. $$F_{holo} = \frac{ZnK_{Zn,mut}}{1+ZnK_{Zn,mut}+\frac{1+ZnK_U}{K_{apo,mut}}}$$

Where $F_{holo}$ is the fraction of folded, zinc-bound DBD, Zn is the concentration of zinc, $K_{Zn,mut}$ is the dissociation constant of a given mutant and $Zn^{2+}$, and $K_U$ is the dissociation constant of unfolded, wild-type DBD and $Zn^{2+}$.

***Cell lines, culture conditions and chemicals.*** TOV112D, H1299, T47D, SUM149P, T98G, Calu-3, U2OS and V138 cells lines were cultured in DMEM with 10% FBS. BXPC3, MCF7 and H460 were cultured in RPMI with 10% FBS. TOV112D, H1299, T47D, SUM149P, T98G, Calu-3, MCF7, U2OS and H460 were purchased from ATCC. V138 was a gift from Dr. Arnold Levine. Cell lines were authenticated by examination of morphology, genotyping by PCR and growth characteristics. The ZMC1 compound was synthesized by Rutgers Molecular Design and Synthesis group, Office of Research and Economic Development [9].

***Generation of expression vectors with various p53 mutants.*** The constructs with various p53 mutants were generated by Site-directed mutagenesis using the Q5 Site-directed mutagenesis kit, following the manufacturer's instructions. The initial plasmid with human *TP53* R273H mutant was purchased from OriGene. The oligos for the mutations are summarized in Table S2.

***Transfection of plasmid constructs and generation of stable cell lines.*** The transfection is performed using Lipofectamine 3000 (Invitrogen), following the manufacturer's instructions. The expression of the p53 protein was confirmed by Western blot. The transfected cells were selected with incubation in the G418-containing medium for stable clones. For the generation of stable cell lines, H1299 cells were transfected with a vector encoding various p53 mutants generated by site-directed mutagenesis. Cells at 80-90% confluence were transfected with Lipofectamine 3000 (Invitrogen) according to manufacturer's instructions. Cells were then selected for the G418 resistance. Single positive clones were isolated and stably maintained in G418-containing medium.

***Cell growth inhibition assay.*** Five thousand cells per well were cultured in 96-well plate such that 50% confluence was reached after one day. At this point, serial dilutions of ZMC1 were added and incubation continued for 3 days. Viability was then measured by Calcein AM (Trevigen). Variable temperature incubations were performed as described in the text.

***Immunofluorescent staining.*** Immunofluorescent staining was performed as described previously [10]. In brief, cells were grown on coverslips, followed by various treatments. The coverslips were fixed with 4% paraformaldehyde and then permeabilized with 0.5% Triton-X100. The conformation of the folded and misfolded/unfolded p53 were recognized respectively by the antibodies PAB1620 (1:50) and PAB240 (1:400) with overnight binding. The secondary antibody, goat anti-mouse IgG, was incubated for 40 m. PAB1620 and PAB240 were purchased from EMD Chemicals. The fluorescent staining intensity was quantified using ImageJ software (National Institutes of Health).

***Immunoprecipitation.*** *C*ells were harvested and lysed using RIPA buffer. The lysates were incubated with protein A/G beads with PAB240/PAB1620 or IgG overnight at 4 °C. The IP products were detected by western blot using the pan-53 antibody DO-7. IgG was used as a negative control. The input total lysates were detected by western blot with p53 antibody DO-1 and actin served as the internal loading control. IgG, DO-7, DO-1, actin antibody, and protein A/G beads were purchased from Santa Cruz Biotechnology.

***Luciferase reporter assay.*** The p53 recognition element (p53RE) in the p21 promoter region, constructed in pGL3 vector, was from Dr. Prives' laboratory (Columbia University, New York, NY, USA). It was transfected into the cells in 96-well plate, followed by the treatment of 1 μM ZMC1 for 48 hours. The luciferase reporter assay was then performed using Dual-Glo luciferase assay system (Promega) following the manufacturer's instructions.

***Data Analysis and Presentation.*** Curve fitting was done with SigmaPlot 13.0, GraphPad Prism and KaleidaGraph 4.5. Unless otherwise noted, all numbers presented are mean ± SE. Two-dimensional plots were generated in Sigmaplot, GraphPad Prism, KaleidaGraph, and R[11], with additional annotation in Adobe Illustrator. Three-dimensional plots were generated in R using the 'lattice' package [12].
